# TopoFlow: Evolutionarily Conditioned Flow Matching for Protein Conformational Ensemble Generation

**DOI:** 10.64898/2026.09.22.753441

**Authors:** Xinguang Yang, Xinyue Cui, Xuhui Li, Dongliang Hou, Suhui Wang, Tengyu Xie, Guijun Zhang

**Affiliations:** College of Information Engineering, Zhejiang University of Technology, 288 Liuhe Road, 310023 Hangzhou, China

## Abstract

Proteins function through dynamic conformational ensembles rather than a single static structure, yet accurately modeling these ensembles remains challenging. Here, we introduce TopoFlow, an evolutionarily conditioned flow-matching framework for protein conformational ensemble generation. TopoFlow uses evolutionary representations from multiple sequence alignments (MSAs) to encode global conformational heterogeneity. It combines this with structural variables from a variational autoencoder (VAE) to capture local flexibility. These complementary representations are then adaptively fused through a conditional modulation module to construct unified conditioning features that guide protein conformational ensemble generation. On the ATLAS molecular dynamics (MD) benchmark, TopoFlow improved the Jensen–Shannon metrics for pairwise distances (JS-PwD), radius of gyration (JS-Rg), and time-lagged independent components (JS-TIC) by 10.73%, 5.78% and 7.75%, respectively, relative to BioEmu. Notably, this performance generalizes to experiment-based evaluation, where TopoFlow exhibits strong agreement with experimental observables, including chemical shifts (CS) and small-angle X-ray scattering (SAXS). Ablation analyses further indicated that the evolutionary representations and structural latent variables contributed complementary information to ensemble generation. Our results suggest that TopoFlow, through its integration of evolutionary and structural latent information, offers a promising strategy for generating structurally plausible and conformationally informative protein ensembles.

## Introduction

Proteins are fundamental biomolecules that perform diverse cellular functions. Many proteins exhibit intrinsic flexibility and populate dynamic conformational ensembles that enable functional adaptation and regulation. These conformational ensembles underlie essential biological processes, including molecular recognition, ligand binding, enzymatic catalysis, allosteric regulation and cellular signaling ^[1–3]^. Therefore, accurately characterizing and generating protein conformational ensembles is essential for understanding molecular mechanisms and advancing computational methods in structural biology.

Experimental techniques such as nuclear magnetic resonance (NMR) ^[4]^ and small-angle X-ray scattering (SAXS) ^[5]^ have long been employed to probe protein conformational ensembles, offering valuable insights into structural flexibility and dynamics in solution. However, these methods typically yield ensemble-averaged measurements, obscuring atomic-level heterogeneity. To explore protein dynamics at all-atom resolution, molecular dynamics (MD) simulations have been widely used to generate protein conformational ensembles by sampling molecular motions under molecular mechanics force fields ^[6,7]^. However, molecular force fields are far from perfect ^[6]^, and the requirement for long-timescale simulations makes MD simulations time-consuming and resource-intensive for large-scale conformational ensemble generation. More recently, deep learning-based structure prediction methods have offered new opportunities for efficiently exploring protein conformational diversity. Methods such as AF-Cluster ^[8]^, AFsample2 ^[9]^ and AFsample3 ^[10]^ have expanded conformational sampling by modifying MSA inputs through clustering or perturbation strategies. While these methods have proven effective in recovering functionally relevant conformational states, there remains room for improvement in both their sampling efficiency and predictive accuracy ^[8–10]^.

In recent years, generative models have emerged as important approaches for modeling protein conformational landscapes by learning and sampling conformational distributions. Early studies using VAEs encoded high-dimensional protein structural representations into continuous latent spaces for conformational sampling ^[11,12]^. Subsequently, diffusion models were increasingly adopted for generating realistic samples from learned data distributions. Representative score-based methods, including Str2Str ^[13]^ and DiG ^[14]^, learn score functions over protein conformational distributions and generate conformational ensembles through annealing-inspired sampling. To improve conformational sampling efficiency while maintaining structural fidelity, recent studies have explored flow-matching frameworks. AlphaFlow and ESMFlow ^[15]^ repurpose AlphaFold2 ^[16]^ and ESMFold ^[17]^ within flow-matching frameworks, conditioning ensemble generation on sequence and MSA information or protein language model representations, respectively. In parallel, BioEmu ^[18]^ leverages large-scale MD simulations and experimental stability data to generate protein ensembles at scale.

Despite recent advances, accurate prediction and reliable evaluation of protein conformational ensembles remain challenging. Two fundamental bottlenecks persist. First, current generative models for protein ensembles are predominantly trained on molecular dynamics (MD)-derived conformational ensembles ^[14,15]^. While capturing functionally relevant conformational states and comprehensively sampling the ensemble landscape requires MD simulations over timescales that are computationally prohibitive for most systems of interest, models trained on such data may inherit sampling biases and fail to represent the full conformational diversity essential for protein function ^[6]^. Second, the evaluation of generated ensembles has largely relied on MD-derived metrics, yet MD simulations themselves are subject to force-field accuracies and its sampling ability. To overcome these limitations, we propose that two complementary strategies are essential—incorporating evolutionary information from multiple sequence alignments (MSAs) to inform functionally relevant motions and validating generated ensembles directly against experimentally accessible observables such as NMR chemical shifts (CSs) and SAXS profiles that report on experimentally observed solution-state behavior ^[19]^.

Here, we present TopoFlow, an evolutionarily conditioned flow-matching framework for generating protein conformational ensembles. The framework integrates information about global conformational heterogeneity with learned structural priors that encode local conformational flexibility. To capture distinct coevolutionary information within the full MSA, we partition the MSA into sequence clusters and separately encode each cluster to obtain cluster-specific evolutionary representations. To obtain local conformational flexibility, a pretrained VAE extracts structural latent variables from MD data. Subsequently, we introduce a conditional modulation module. This module uses structural latent variables to adaptively modulate the evolutionary representations, thereby conditioning the flow-matching model to generate protein conformations from noisy rigid frames. Across benchmarks based on MD data ^[7]^, experimental observables ^[19]^ and alternative conformations ^[18]^, TopoFlow outperforms or remains competitive with representative baseline methods, demonstrating its potential to generate structurally plausible and conformationally informative protein ensembles.

## Results

### Overview of TopoFlow

TopoFlow generates conformational ensembles by combining evolutionary representations, structural latent variables and flow-matching generation (**Fig. 1a**). Given an input sequence, an MSA is first searched and partitioned into clusters ^[8]^, with each cluster processed by a frozen MSA encoder ^[16]^ to obtain cluster-specific single and pair representations. In parallel, a VAE pretrained on MD-derived structural features provides structural latent variables that encode local conformational priors ^[20]^. The Conditional Modulation module integrates these cluster-specific single and pair representations and structural latent variables to construct conditioning features for the Flow Matching module (**Fig. 1b**). These conditions guide the Flow Matching module ^[21]^ (**Fig. 1c**), which iteratively updates noised backbone frames to generate conformational ensembles.

**Fig. 1.**
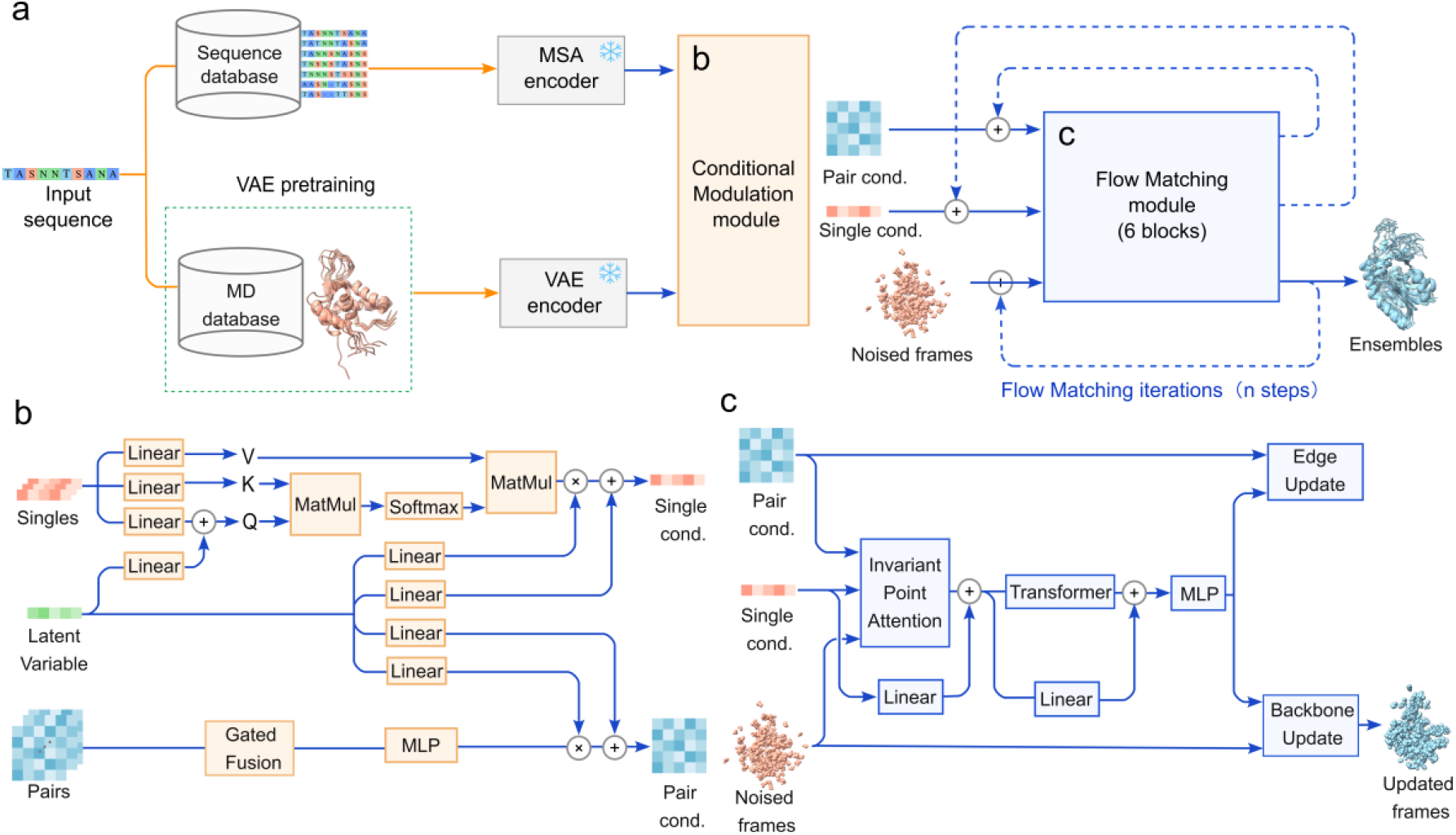
Architecture of the TopoFlow framework. **a**, Overview of TopoFlow pipeline. **b**, Conditional Modulation module. **c**, Flow Matching module.

### Performance on MD simulation benchmarks

We first evaluated TopoFlow on the ATLAS test set to determine whether the generated ensembles reproduce the conformational distributions sampled by the reference MD trajectories (**Fig. 2**). The comparison included BioEmu ^[18]^, AlphaFlow ^[15]^, Str2Str ^[13]^, and idpGAN ^[22]^. For each target protein, TopoFlow and the competing methods generated 1,000 conformations, and the generated ensembles were compared with the corresponding reference MD ensembles using complementary metrics, including JS-PwD, JS-Rg, JS-TIC, RMSF correlation, weak and transient contacts, with Jaccard similarity used to quantify the agreement between generated and reference contact sets.

**Fig. 2.**
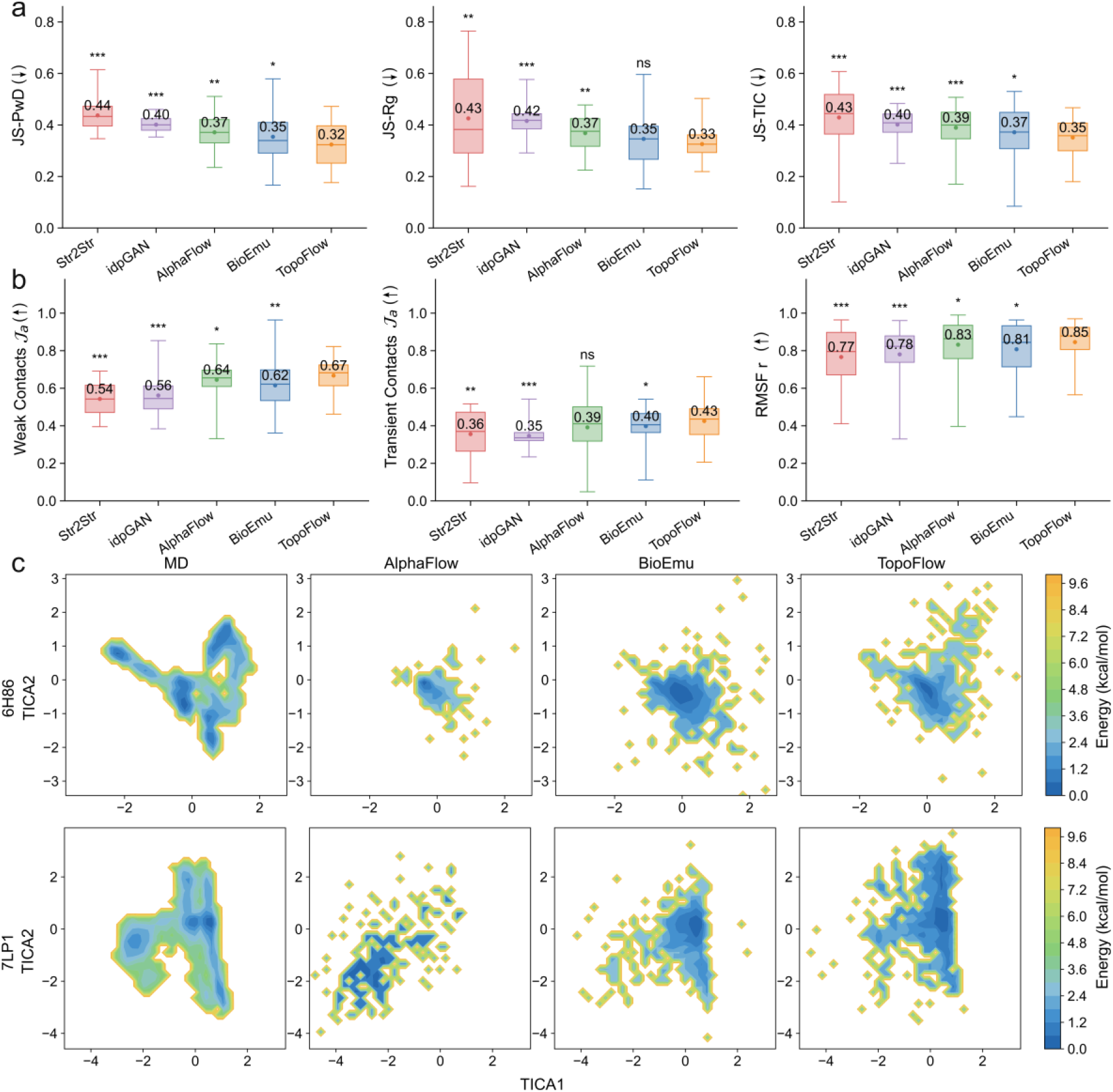
Performance on MD simulation benchmarks. **a**, Distributional agreement measured by JS-PwD, JS-Rg and JS-TIC. **b**, Weak contacts *Ja*, transient contacts *Ja* and Pearson correlation of RMSF (RMSF r). **c**, Representative TICA landscapes for 6H86 and 7LP1 comparing BioEmu, AlphaFlow and TopoFlow with reference MD ensembles. The color scale represents the relative conformational free energy landscape estimated from the sampling density in the TICA space, where lower-energy regions correspond to highly populated conformational states. Box plots show the median and interquartile range (IQR), with the central line indicating the median and the box representing the 25th–75th percentiles. Numbers inside the boxes indicate mean values. Arrows indicate the preferred direction of each metric, with ↑ and ↓ denoting higher and lower values are better, respectively. Statistical significance was assessed using Wilcoxon signed-rank tests ^[25]^. ns, not significant; \**P* < 0.05; \*\**P* < 0.01; \*\*\**P* < 0.001.

As shown in **Fig. 2a**, TopoFlow achieved improved agreement with reference MD ensembles on the ATLAS test set. TopoFlow improved JS-PwD, JS-Rg, and JS-TIC by 10.73%, 5.78%, and 7.75%, respectively, relative to the best competing method (**Table S1**). Statistical analysis showed TopoFlow significantly outperformed Str2Str, idpGAN, and AlphaFlow across the distributional metrics. We further assessed whether TopoFlow reproduced dynamic properties captured by reference MD ensembles (**Fig. 2b**). Relative to the best-performing baseline, TopoFlow improved Jaccard similarities (*Ja*) of weak contacts, the *Ja* of transient contacts, and RMSF correlation by 3.57%, 6.78%, and 3.02%, respectively. These improvements were statistically significant, with TopoFlow outperforming all four baselines in weak contacts *Ja* and RMSF correlation. Statistical analysis showed that TopoFlow significantly outperformed Str2Str, idpGAN, and BioEmu for transient contacts *Ja*, while exhibiting comparable performance to AlphaFlow. Detailed performance comparisons are provided in **Supplementary Table S1**. These results demonstrate that TopoFlow effectively recovers MD-derived conformational distributions while capturing dynamic residue interactions and local flexibility underlying protein conformational motions.

To further evaluate the similarity of conformational landscapes generated by TopoFlow and baseline methods in the time-lagged independent component analysis (TICA) ^[18]^ space, we selected two representative targets (PDB IDs:6H86 ^[23]^ and 7LP1 ^[24]^) (**Fig. 2c**). 6H86 corresponds to the mouse synaptonemal complex central element protein SYCE3, which is involved in meiotic chromosome synapsis and synaptonemal complex assembly, whereas 7LP1 is an NEDD4L WW domain involved in protein–protein recognition through PPxY motif binding. All generated conformations were projected onto the first two TICA components derived from the corresponding reference MD ensembles. Compared with baseline methods, TopoFlow-generated conformations better recover the reference MD ensembles.

### Performance on experimental observables

We next evaluated whether TopoFlow generated experimentally consistent conformational ensembles on the PeptoneDB-CS and PeptoneDB-SAXS benchmarks (**Fig. 3**,**4**). For each target, 100 conformations were generated for ensemble-level evaluation. We analyzed proteins in the ordered regime (mean G-score < 0.5) ^[19]^ as the primary evaluation, while extending the analysis to the full G-score range in **Supplementary Fig. S3 and S4**. The corresponding experimental observables were calculated from the generated ensembles using the corresponding forward models ^[19,26,27]^, where chemical shifts were used to assess local secondary structure content and SAXS profiles were used to characterize the overall shape of the protein ensemble. Following the PeptoneBench ^[19]^ evaluation protocol, agreement between predicted and experimental observables was quantified using uncertainty-normalized RMSE before and after maximum entropy reweighting and summarized using LOWESS-AUC across different G-score ranges.

**Fig. 3.**
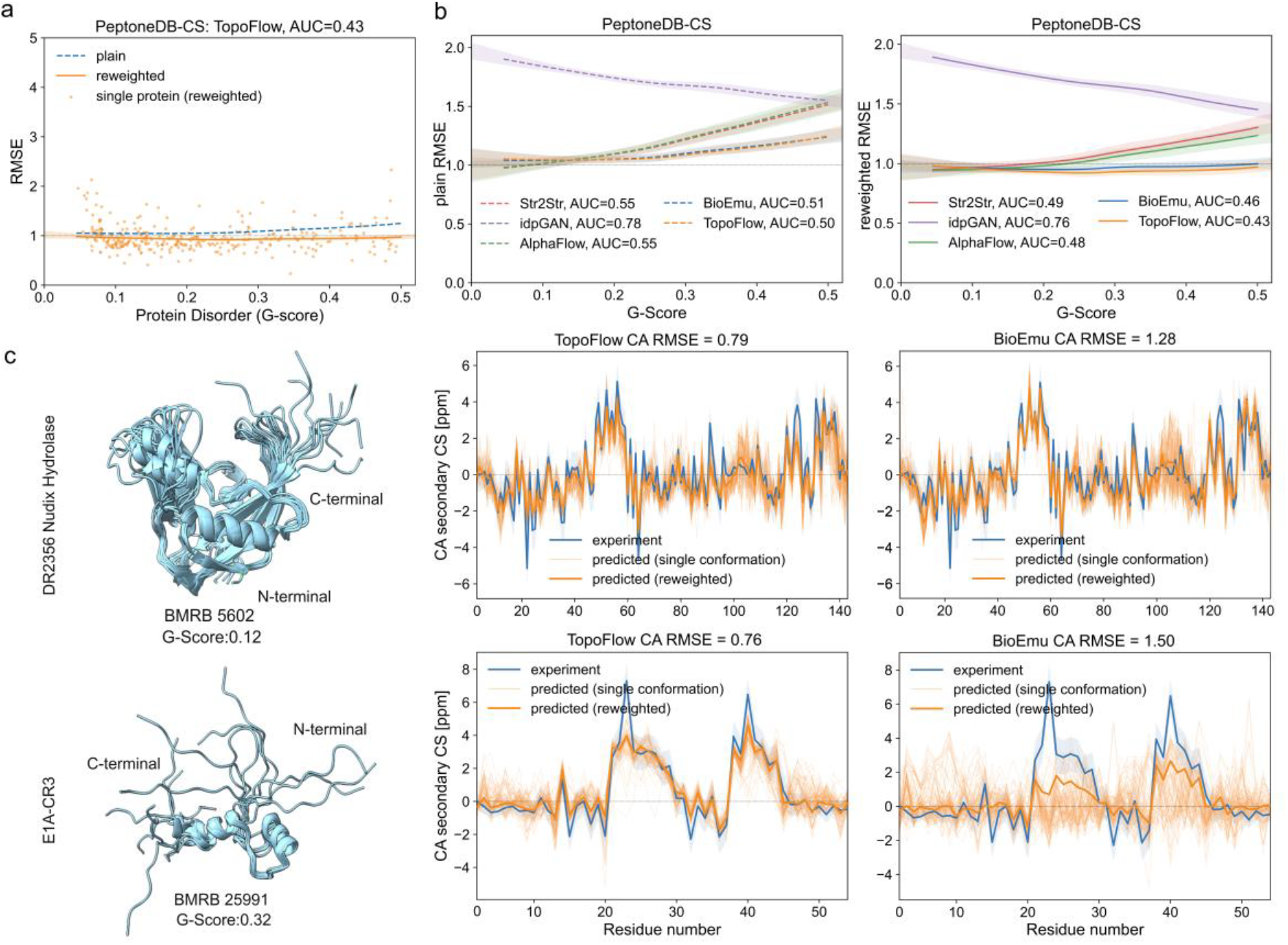
Agreement with PeptoneDB-CS chemical shift observables. **a**, Normalized RMSE of TopoFlow as a function of mean G-score before and after maximum entropy reweighting. Points represent individual proteins, and the dashed and solid curves indicate LOWESS fits before and after reweighting, respectively ^[19]^. **b**, LOWESS-smoothed normalized RMSE curves comparing TopoFlow with baseline methods before and after ensemble reweighting across the G-score range. **c**, Representative chemical shift cases for BMRB 5602 and BMRB 25991. The corresponding generated ensembles and G-score values are shown on the left. Experimental CA secondary chemical shifts are compared with predictions from individual generated conformations and the reweighted ensemble average for TopoFlow and BioEmu, with normalized RMSE values shown above each plot.

**Fig. 4.**
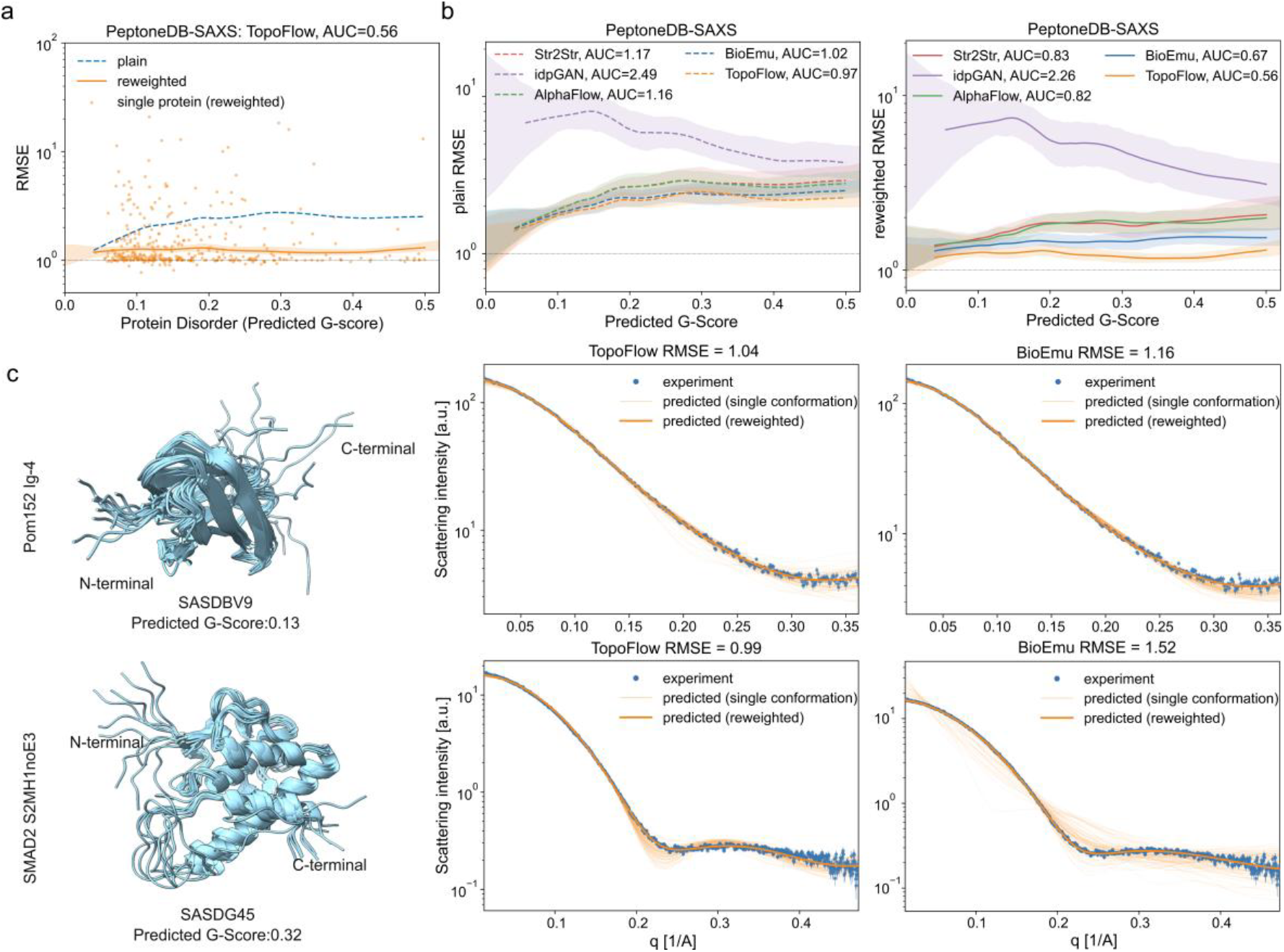
Agreement with PeptoneDB-SAXS observables. **a**, Normalized RMSE of TopoFlow as a function of predicted G-score before and after maximum entropy reweighting. Points represent individual proteins, and the dashed and solid curves indicate LOWESS fits before and after reweighting, respectively ^[19]^. **b**, LOWESS-smoothed normalized RMSE curves comparing TopoFlow with baseline methods before and after ensemble reweighting across the predicted G-score range. **c**, Representative SAXS cases for SASDBV9 and SASDG45. The corresponding generated ensembles and predicted G-score values are shown on the left. Experimental SAXS profiles are compared with predictions from individual generated conformations and the reweighted ensemble average for TopoFlow and BioEmu, with normalized RMSE values shown above each plot.

As shown in **Fig. 3a**, chemical shifts were calculated from TopoFlow-generated ensembles on the PeptoneDB-CS benchmark. The generated ensemble was first evaluated directly using the original conformational populations (referred to as plain). Subsequently, maximum entropy reweighting was applied to adjust conformational populations while incorporating experimental chemical shift information, generating a reweighted result. The ensemble-averaged chemical shifts from both approaches were compared with experimental measurements, with the reweighted ensemble showing improved agreement. Maximum entropy reweighting generally reduced the uncertainty-normalized RMSE for proteins in the ordered regime, decreasing the LOWESS-AUC from 0.50 to 0.43. As shown in **Fig. 3b**, the reweighted TopoFlow ensemble achieved a LOWESS-AUC comparable to that of BioEmu (0.46) and lower than those of AlphaFlow (0.48), Str2Str (0.49), and idpGAN (0.76). Notably, TopoFlow and BioEmu exhibited different performance trends across disorder ranges. TopoFlow achieved better agreement with experimental chemical shifts in the ordered regime, whereas BioEmu maintained more consistent performance across the full G-score range (**Fig. S3 and Table S2**).

We further illustrated the performance of TopoFlow using two representative chemical shift cases with low G-scores (**Fig. 3c**). BMRB 5602 (G-score 0.12) corresponds to Nudix hydrolase DR2356 (DR2356 Nudix Hydrolase) from Deinococcus radiodurans ^[28]^, a nucleotide-metabolizing enzyme from an extremely stress-resistant bacterium. For this target, TopoFlow closely matched the experimental CA secondary chemical shifts and achieved a CA RMSE of 0.79, compared with 1.28 for BioEmu. BMRB 25991 (G-score 0.32) corresponds to the CR3 region of the human adenovirus E1A protein (E1A-CR3) ^[29]^, a viral regulatory region involved in transcriptional activation. TopoFlow also captured the main residue-level chemical shift features and achieved a CA RMSE of 0.76, compared with 1.50 for BioEmu. Together, these examples highlight the ability of TopoFlow to achieve high consistency with experimental chemical shifts for proteins with low G-scores.

We next evaluated TopoFlow on the PeptoneDB-SAXS benchmark to assess the agreement between predicted and experimental SAXS profiles (**Fig. 4**). Maximum entropy reweighting generally reduced the normalized RMSE for proteins in the ordered regime, decreasing the LOWESS-AUC from 0.97 to 0.56 (**Fig. 4a**). As shown in **Fig. 4b**, the reweighted TopoFlow ensemble showed improved performance compared with BioEmu (0.67), AlphaFlow (0.82), Str2Str (0.83), and idpGAN (2.26). Analysis across the G-score range showed that TopoFlow maintained competitive agreement with SAXS observables across the ordered regime, while higher RMSE values were mainly observed for highly disordered proteins (**Fig. S4 and Table S3**).

To illustrate the ability of TopoFlow to capture experimental scattering behavior for proteins with distinct structural characteristics, we examined representative SAXS cases (**Fig. 4c**). SASDBV9 (predicted G-score 0.13) corresponds to an immunoglobulin-like domain of the yeast nucleoporin Pom152 (Pom152 Ig-4) ^[30]^, whereas SASDG45 (predicted G-score 0.32) corresponds to an N-terminal MH1 domain construct of human SMAD2 (SMAD2 S2MH1noE3) ^[31]^. Compared with BioEmu, TopoFlow-generated ensembles showed closer agreement with experimental SAXS profiles after reweighting for both targets. The reweighted SAXS profiles calculated from TopoFlow ensembles closely followed the experimental profiles across the measured range of the scattering-vector magnitude. This agreement indicates that the generated ensembles captured the overall shape of the protein ensemble.

### Performance on alternative conformations

We evaluated the ability of TopoFlow to capture multiple functionally relevant conformational states using the domain motion benchmark. In addition to generative model-based approaches, we further included AF-Cluster ^[8]^ and AFsample3 ^[10]^, two recently developed methods specifically designed to generate alternative conformational states, as representative benchmarks for functional conformational sampling. For each protein target, each method generated 1000 conformations, and the best-matching structures were selected for each reference state using RMSD and TM-score ^[32]^.

As shown in **Fig. 5a**, TopoFlow generated conformations that closely matched experimentally resolved reference states, achieving RMSD values of 1.82 Å and 1.68 Å and TM-scores of 0.92 and 0.94 for closed and open states, respectively (**Table S2**). These results were comparable to those of representative baseline methods, including BioEmu and AFsample3. To quantify the recovery of both reference states, we assessed whether each reference state was recovered using an RMSD threshold of ≤3 Å. TopoFlow successfully recovered 18 of 22 closed-state references (82%) and 18 of 22 open-state references (82%) (**Fig. S5 and Table S5**). Together, these results demonstrate the potential of TopoFlow for recovering functionally relevant conformational states.

**Fig. 5.**
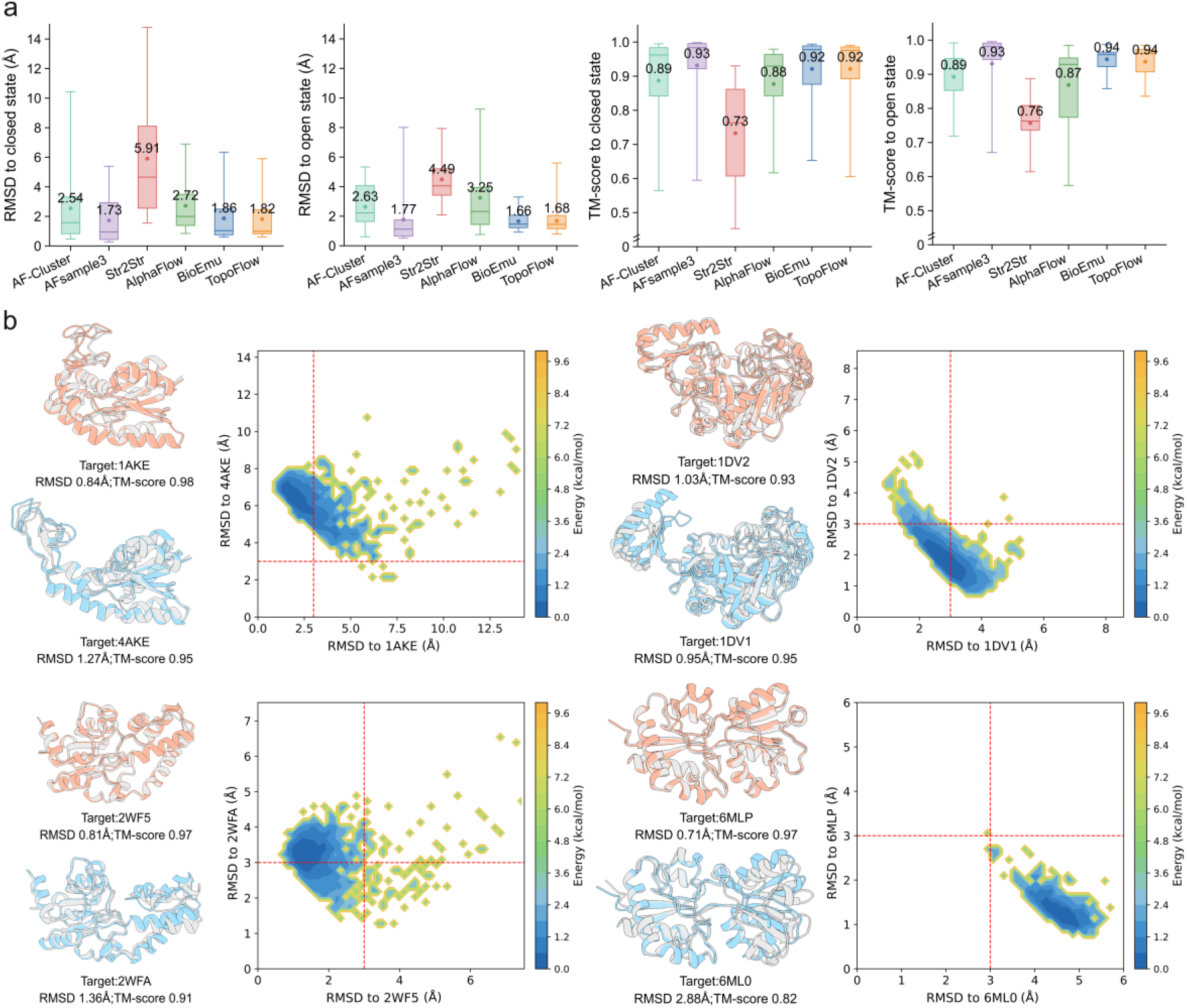
Performance on alternative conformations. **a**, Distributions of global RMSD and TM-score values calculated from the same best-matched generated conformations across all domain motion targets. **b**, RMSD-to-state scatter plots for four representative proteins. Red dashed lines indicate the 3 Å RMSD recovery threshold; points below the horizontal line or to the left of the vertical line are within 3 Å of the corresponding reference state. The color scale represents the conformational sampling density in the RMSD space relative to the two reference states, with higher-density regions indicating frequently sampled conformational states.

**Fig. 5b** shows representative RMSD scatter plots relative to the two experimentally resolved reference states. The four targets span distinct types of large-scale domain motion, including the open–closed transition of adenylate kinase during nucleotide phosphotransfer ^[33]^, conformational rearrangement of biotin carboxylase during biotin carboxylation ^[34]^, catalytic-domain motion of β-phosphoglucomutase during phosphate transfer ^[35]^, and ligand-induced opening and closing of the LAO periplasmic binding protein ^[36]^. In these representative cases, TopoFlow sampled diverse conformations distributed near both reference states, indicating that the generated ensembles capture large-scale functional domain motions.

### Ablation Study

To evaluate the individual and combined contributions of the two key generation conditions in TopoFlow, we performed ablation on the ATLAS test set (**Fig. 6**). The full model (TopoFlow) was compared with three ablated variants, including a model without the VAE-derived structural latent variables (w/o VAE), a model in which cluster-specific MSA conditioning was replaced by conditioning derived from the original unclustered MSA (w/o Cluster), and a model incorporating both modifications simultaneously (w/o Cluster & VAE).

**Fig. 6.**
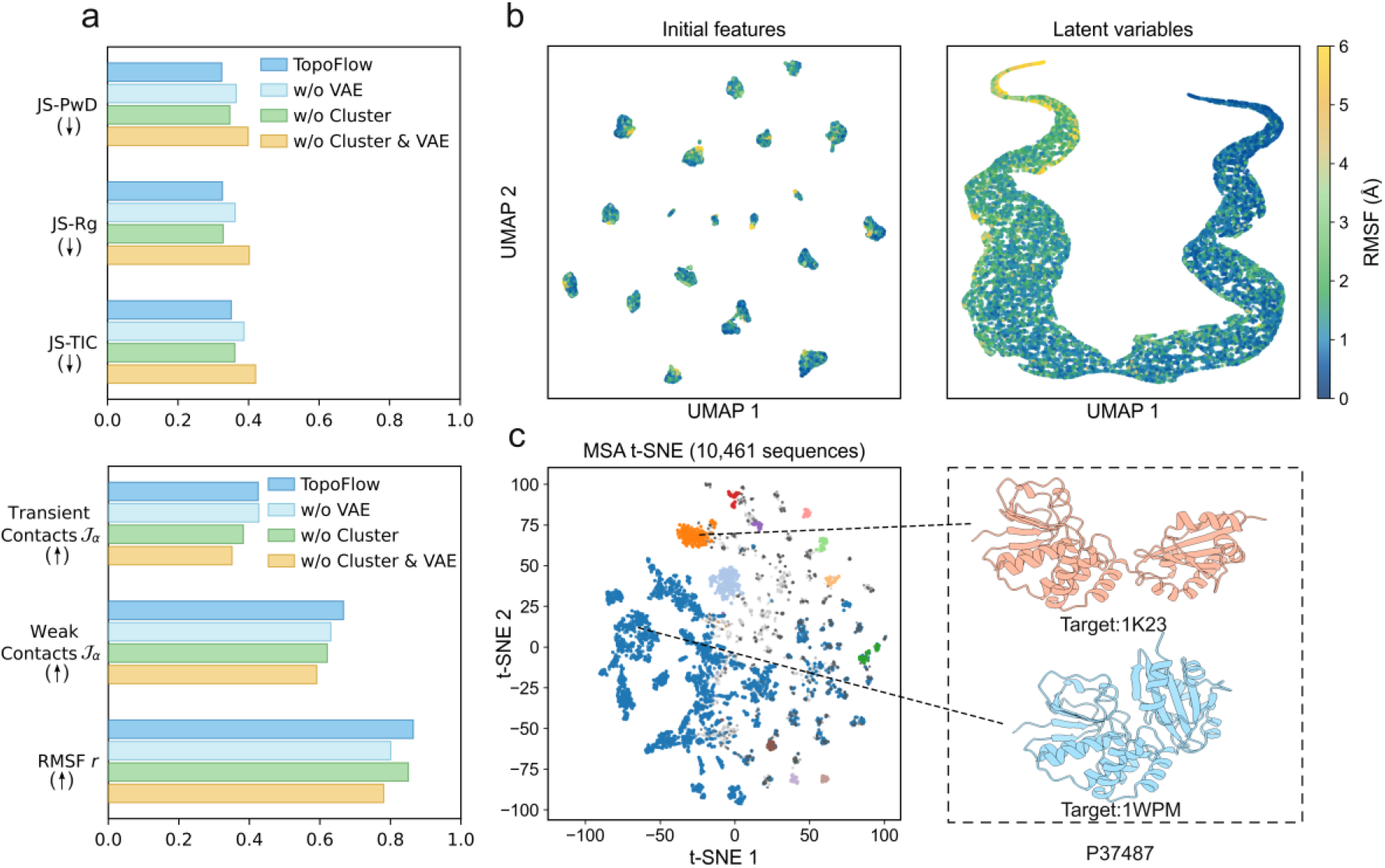
Ablation and explainability analyses. **a**, Ablation analysis of structural latent variables from the VAE encoder and MSA clustering across MD benchmark metrics. **b**, UMAP projections of initial structural features and VAE encoder latent variables coloured by RMSF. **c**, t-SNE projection of the clustered MSA.

The full TopoFlow model achieved the best or near-best performance across the evaluated metrics, including the lowest JS-PwD, JS-Rg, and JS-TIC values (**Fig. 6a**). Removing the VAE-derived structural latent variables mainly affected the distributional metrics and RMSF correlation (**Table S6**), suggesting that these structural latent variables contribute to distributional accuracy and residue-level flexibility. In contrast, replacing clustered MSA conditions with unclustered MSA had a stronger effect on weak and transient contact agreement (**Table S6**), suggesting that clustered MSA are particularly useful for capturing weak and transient contacts within generated ensembles. The variant combining both modifications showed the largest overall decline, supporting the complementary contributions of VAE-derived structural latent variables and clustered MSA to ensemble generation.

To further explain these ablation results, we analyzed the representations used by TopoFlow. UMAP ^[37]^ projections showed that VAE-derived structural latent variables provided a more continuous low-dimensional representation of conformational features than the initial structural features and showed a clearer correspondence with residue-level RMSF (**Fig. 6b**). In parallel, for the representative target Inorganic Pyrophosphatase (P37487), MSA clustering produced distinct sequence groups in the t-SNE embedding, indicating that clustered MSA conditions capture sequence diversity within the MSA representation (**Fig. 6c**). Together, these analyses provide additional support for the ablation results by showing the organization of structural clustered MSA representations and latent variables.

## Conclusions

In this study, we introduced TopoFlow, an evolutionarily conditioned flow-matching framework for protein conformational ensemble generation. TopoFlow leverages clustered MSA-derived evolutionary representations to capture heterogeneous evolutionary constraints and combines them with VAE-derived structural latent variables that encode local conformational flexibility. These complementary representations are integrated through a Conditional Modulation module to guide conformational sampling within the flow-matching framework. Across the ATLAS ^[7]^ MD benchmark, the PeptoneDB-CS and PeptoneDB-SAXS benchmarks ^[19]^ based on experimental observables, and the domain motion benchmark ^[18]^, TopoFlow achieved competitive or improved performance in reproducing conformational distributions, residue-level flexibility, weak and transient contacts, experimental observables, and alternative conformational states. These results support the idea that heterogeneous evolutionary information and learned structural flexibility provide complementary constraints for modeling protein conformational ensembles.

Despite these encouraging results, TopoFlow still has room for improvement in applicability and accuracy. TopoFlow showed reduced performance for highly disordered proteins, whose broad, heterogeneous conformational landscapes are harder to characterize and sample accurately. In addition, experimental observables are currently used primarily for evaluating the generated ensembles rather than as direct constraints during model training or conformational generation. Future work could therefore focus on improving the sampling of highly disordered proteins and incorporating experimental information, such as chemical shifts and SAXS measurements, into the training or generation process to better constrain solution-state conformational ensembles. These developments could further improve the accuracy and applicability of TopoFlow across a broader range of proteins.

## Methods

### Data Set

Training data consisted of experimentally determined structures from the PDB and conformational snapshots generated from ATLAS ^[7]^ MD simulations. For the PDB training set, we selected monomeric protein structures from PDB entries released before 9 June 2023 and filtered them based on experimental resolution (<2.5 Å) and sequence length (10–512 residues). To capture the structural diversity represented by homologous proteins, we clustered the remaining protein sequences using MMseqs2 ^[38]^ with a 40% sequence identity threshold. We further removed clusters containing proteins sharing more than 40% sequence identity with any of the test datasets to prevent information leakage. The final PDB training set comprised 4,686 sequence clusters containing 26,193 monomeric protein structures. Structures are sampled randomly from these clusters at each training iteration. For the ATLAS training set, proteins were selected according to the temporal split strategy used in AlphaFlow ^[15]^ and further filtered to remove redundancy with proteins in all test datasets, resulting in 1,238 non-redundant proteins. For each retained protein, 100 conformational snapshots were sampled from three 100 ns trajectories, yielding 123,800 structures in total.

Test data included the ATLAS test set ^[15]^, PeptoneDB-CS ^[19]^, PeptoneDB-SAXS ^[19]^, and the domain motion benchmark introduced by BioEmu ^[18]^. The ATLAS test set was used to evaluate agreement with MD-derived metrics and consists of 82 protein systems from standardized all-atom MD trajectories following the temporal split introduced by AlphaFlow ^[15]^. PeptoneDB-CS and PeptoneDB-SAXS were used to evaluate consistency with solution-state experimental observables ^[19]^. PeptoneDB-CS contains 659 protein entries with NMR chemical shift data curated from the Biological Magnetic Resonance Data Bank (BMRB) ^[39]^, whereas PeptoneDB-SAXS contains 439 protein entries with SAXS profiles curated from SASBDB ^[40]^. The domain motion benchmark was used to assess recovery of alternative functional conformations, comprising 22 protein targets with large-scale conformational changes and two experimentally resolved reference states for each target. The open and closed states were defined based on differences in the radius of gyration (Rg) between the two reference conformations, with detailed information provided in **Supplementary Table S7**.

### Evolutionary Information Embedder

For each input sequence, MSAs were generated with ColabFold ^[41]^ using MMseqs2 ^[38]^ searches against UniRef30 ^[42]^, BFD ^[43]^ and MGnify ^[44]^. UniRef30 provides a redundancy-reduced collection of clustered UniProt sequences ^[42]^, BFD contributes large-scale sequence diversity from genomic and metagenomic resources ^[43]^, and MGnify expands coverage with protein sequences predicted from environmental and microbiome datasets ^[44]^. These complementary databases were used to increase MSA depth and diversity. The resulting MSA sequences were clustered using DBSCAN ^[8]^, and five representative evolutionary clusters were selected according to the clustering procedure described in **Supplementary Note S1**.

For each of the five clustered MSAs, residue-level single and pair representations were extracted using a frozen MSA encoder composed of five independent Evoformer modules adapted from AlphaFold2 ^[16]^. Each Evoformer module contains 48 blocks and processes one MSA cluster independently. The resulting cluster-specific single and pair representations were used as fixed conditions during both training and inference, providing global heterogeneous evolutionary context for conformational ensemble generation.

### Local Conformation Embedder

To capture local conformational flexibility from structural information, we pretrained a VAE using MD conformational training set. For each sampled structure, Rosetta one-body and two-body energy terms ^[45]^ were computed to characterize local energetic environments and pairwise interactions, while residue-pair distance and orientation maps were derived from backbone geometry to encode spatial relationships between residues. These energetic and geometric features were jointly used as structural inputs for VAE pretraining, where the encoder predicts the parameters mean (*µ*) and standard deviation (*σ*) of residue-wise latent distributions to learn latent representations of local conformational flexibility. The VAE consists of an equivariant graph neural network (EGNN) ^[46]^ and Transformer ^[47]^ blocks and predicts a residue-wise latent distribution from the input sequence. Detailed descriptions of the model architecture and pretraining procedures are provided in **Supplementary Note S2, Fig. S1**, and **Supplementary Algorithm S1**.

For each input sequence, the pretrained VAE encoder is used to obtain a residue-wise Gaussian latent prior parameterized by *µ*_*ϕ,i*_ and *σ*_*ϕ,i*_, from which structural latent variables are sampled using the reparameterization strategy ^[48]^:

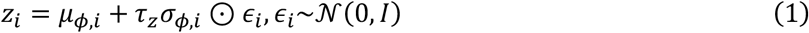

where *µ*_*ϕ,i*_ and *σ*_*ϕ,i*_ denote the latent mean and standard deviation for residue *i, ϵ*_*i*_ is sampled from the standard Gaussian distribution *N*(0, *I*), ⊙ denotes element-wise multiplication, and *τ*_*z*_ controls the latent sampling temperature. The sampled latent variables encode local conformational flexibility and are subsequently used by the Conditional Modulation module to construct conditioning features for flow matching, remaining fixed throughout the generation process.

### Conditional Modulation module

The Conditional Modulation module adaptively integrates cluster-specific evolutionary representations from the MSA encoder, providing global conformational heterogeneity, with structural latent variables from the VAE encoder, capturing local conformational flexibility. The fused conditioning features are then used to guide the Flow Matching module for ensemble generation. Specifically, the unified conditioning features are constructed through latent-conditioned attention over evolutionary representations. For the single condition, cluster-specific single representations are first projected into query, key and value features. Structural latent variables are used to generate latent-dependent attention biases that modulate the aggregation of evolutionary information across MSA clusters. For residue *i*, the query feature 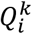 is derived from the single representation of MSA cluster *k*, whereas the key and value features 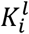 and 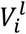 are derived from the single representation of MSA cluster *l*. The latent-conditioned attention weight is computed as:

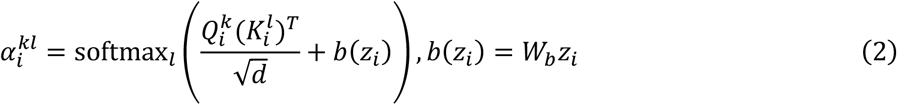

where 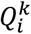 and 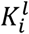 denote the query and key features derived from the single representations of MSA clusters *k*and *l*, respectively; *d* is the feature dimension, and *b*(*z*_*i*_) is the latent-dependent attention bias obtained through a learnable linear projection *W*_*b*_.

Following aggregation over the key–value cluster index *l* according to the latent-conditioned attention weights, the resulting representations are further modulated by latent-dependent scale and bias terms derived from the structural latent variables to generate the cluster-specific single condition:

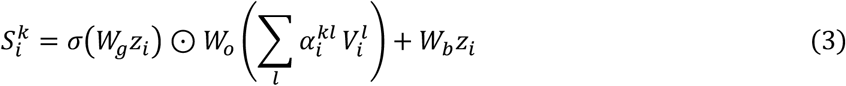

where 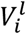 denotes the value feature derived from the single representation of MSA cluster *l*; *W*_*g*_, *W*_*o*_, and *W*_*b*_ are learnable projection matrices; and 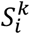 represents the single condition generated for query cluster *k*. The cluster-specific single conditions are then aggregated across query clusters to obtain the final single condition *S*_*i*_.

In parallel, the pair branch independently applies latent-conditioned gating and modulation to pair representations to generate the pair condition:

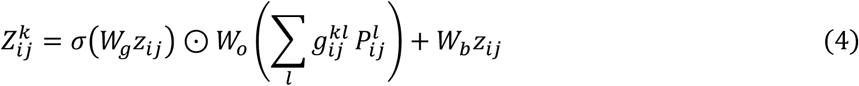

where 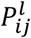 denotes the pair representation between residues *i* and *j* derived from the pair representation of MSA cluster *l, z*_*ij*_ represents the pairwise structural latent variable, 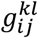 denotes the gating weight between query MSA cluster *k* and key-value MSA cluster *l*, and *σ*(⋅) denotes the sigmoid function. The latent-conditioned gating first aggregates pair representations across MSA clusters, after which the aggregated features are modulated by latent-dependent scale and bias terms to generate the cluster-specific pair condition 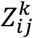. The resulting pair representations constitute the pair condition *Z*_*ij*_. Together, the single and pair conditions generated by the Conditional Modulation module are subsequently provided to the Flow Matching module.

### Flow Matching module

The Flow Matching module predicts backbone-frame updates under the guidance of the single and pair conditions generated by the Conditional Modulation module. At time *t*, it receives the current noised backbone frames *T*_*t*_, the single condition *S*_*i*_ and the pair condition *Z*_*ij*_ (**Fig. 1c**). The module contains six recycled update blocks that iteratively update the backbone frames while incorporating these conditioning features. Each block consists of invariant point attention, Transformer layers, an edge update module and a backbone update module. Invariant point attention ^[16]^ incorporates spatial context from the current backbone frames while preserving equivariance to global rigid-body transformations, whereas the Transformer layers capture residue-level contextual dependencies conditioned on the structural information provided by the current backbone frames. Through recycling, the six update blocks are repeatedly applied to iteratively refine the geometry of backbone frames. During training, the Flow Matching module is optimized using the conditional flow-matching objective. During inference, the conditional probability flow is modeled by an ordinary differential equation (ODE) ^[49]^, which is discretized using linear interpolation for conformational sampling. The sampling trajectory is divided into 100 integration steps with a constant interval of Δ*t* = 0.01. A single integration step is given by ^[50]^:

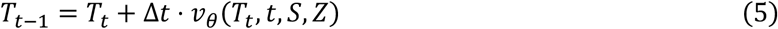

where *T*_*t*_ denotes the frames at time *t*, and *v*_*θ*_(*T*_*t*_, *t, S, Z*) denotes the learned conditional vector field that determines the update direction of the conformational state. Detailed training and inference procedures are provided in **Supplementary Note S3** and **Algorithms S2–S3**.

### Evaluation metrics

For the ATLAS test set, we used JS-PwD and JS-Rg to evaluate the consistency of generated ensembles with reference MD ensembles in terms of global structural distributions and protein compactness, respectively ^[13]^. JS-TIC was used to assess dominant conformational motions from the reference MD trajectories ^[18]^. Weak and transient contacts were used to characterize dynamic residue–residue contact dynamics during conformational sampling, and Jaccard similarity (*Ja*) was calculated to quantify the agreement between generated and reference contact sets ^[15]^. Finally, RMSF correlation was calculated to assess the consistency of residue-level flexibility between generated and reference ensembles ^[15]^.

For the PeptoneDB-CS and PeptoneDB-SAXS benchmarks, we used uncertainty-normalized root mean square error (RMSE) before and after maximum entropy reweighting to quantify the agreement between generated conformational ensembles and experimental measurements ^[19]^. In addition, the area under the LOWESS curve (LOWESS-AUC) was used to summarize the overall performance across different G-score ^[19]^ ranges.

For the domain motion benchmark introduced by BioEmu ^[18]^, we used global root-mean-square deviation (RMSD) and TM-score ^[32]^ to evaluate the recovery of experimentally resolved alternative conformational states. RMSD was used to quantify structural deviation from each reference conformation, whereas TM-score was used to assess overall structural similarity ^[32]^. Following the evaluation protocol used in BioEmu, a reference state was considered recovered if at least 0.1% of the generated conformations achieved an RMSD of ≤3 Å to that state ^[18]^. Detailed definitions of all metrics are provided in **Supplementary Note S4**.

## Data and Software Availability

The authors declare that the data supporting the findings and conclusions of this study are available within the paper and its Supporting Information. Public benchmark datasets used in this work were obtained from the previously published PeptoneBench ^[13]^, ATLAS ^[15]^, and the domain motion benchmark from BioEmu ^[18]^ resources. The processed benchmark inputs, evaluation results, and generated conformational ensemble summaries are available in the TopoFlow repository upon publication. The TopoFlow source code and inference scripts are publicly available at https://github.com/iobio-zjut/Topoflow. The TopoFlow web server is freely accessible at http://zhanglab-bioinf.com/TopoFlow/.

All comparative methods were evaluated using publicly available implementations released by the original authors. For MD-based ensemble evaluation, BioEmu ^[18]^, AlphaFlow ^[15]^, Str2Str ^[13]^, and idpGAN ^[22]^ were used as baseline methods. For alternative conformational state recovery, AF-Cluster ^[8]^and AFsample3 ^[10]^ were included as additional baseline methods. The recommended settings from the corresponding publications were adopted whenever applicable.

## Author contributions

G.Z. conceived and supervised the research. G.Z., X.Y., X.C., and X.L. designed the study. X.Y., X.C., X.L., D.H., and S.W. developed the methodology, performed computational analyses, and collected the data. G.Z., X.Y., X.C., X.L., and D.H. analyzed and interpreted the results. G.Z., X.Y. and X.C. wrote the manuscript. G.Z. and T.X. revised the manuscript. All authors discussed the results and approved the final version of the manuscript.

## Acknowledgments

This work was supported by the National Key R&D Program of China (2022ZD0115103), the National Nature Science Foundation of China (62203389), the “Pioneer” and “Leading Goose” R&D Program of Zhejiang (2025C01190), the Zhejiang Province High-level Talent Special Support Program (2023R5248).

## Competing interests

The authors declare no competing interests.

## Additional information

Correspondence and requests for materials should be addressed to G.Z.

## Notes

### Competing Interest Statement

The authors have declared no competing interest.

## REFERENCES

1. G. Wei, W. Xi, R. Nussinov, and B. Ma, “Protein Ensembles: How Does Nature Harness Thermodynamic Fluctuations for Life? The Diverse Functional Roles of Conformational Ensembles in the Cell,” Chemical Reviews 116, no. 11 (2016): 6516–6551, 10.1021/acs.chemrev.5b00562.

2. T. Xie, T. Saleh, P. Rossi, and C. G. Kalodimos, “Conformational States Dynamically Populated by a Kinase Determine Its Function,” Science 370, no. 6513 (2020): eabc2754, 10.1126/science.abc2754.

3. X. Cui, L. Ge, X. Chen, et al., “Beyond Static Structures: Protein Dynamic Conformations Modeling in the Post-AlphaFold Era,” Briefings in Bioinformatics 26, no. 4 (2025): bbaf340, 10.1093/bib/bbaf340.

4. R. Zadorozhnyi, A. M. Gronenborn, and T. Polenova, “Integrative Approaches for Characterizing Protein Dynamics: NMR, CryoEM, and Computer Simulations,” Current Opinion in Structural Biology 84 (2024): 102736, 10.1016/j.sbi.2023.102736.

5. M. L. Mugnai, D. Chakraborty, H. T. Nguyen, et al., “Sizes, Conformational Fluctuations, and SAXS Profiles for Intrinsically Disordered Proteins,” Protein Science 34, no. 4 (2025): e70067, 10.1002/pro.70067.

6. K. Lindorff-Larsen, S. Piana, R. O. Dror, and D. E. Shaw, “How Fast-Folding Proteins Fold,” Science 334, no. 6055 (2011): 517–520, 10.1126/science.1208351.

7. Y. Vander Meersche, G. Cretin, A. Gheeraert, J.-C. Gelly, and T. Galochkina, “ATLAS: Protein Flexibility Description from Atomistic Molecular Dynamics Simulations,” Nucleic Acids Research 52, no. D1 (2024): D384–D392, 10.1093/nar/gkad1084.

8. H. K. Wayment-Steele, A. Ojoawo, R. Otten, et al., “Predicting Multiple Conformations via Sequence Clustering and AlphaFold2,” Nature 625, no. 7996 (2024): 832–839, 10.1038/s41586-023-06832-9.

9. Y. Kalakoti, and B. Wallner, “AFsample2 Predicts Multiple Conformations and Ensembles with AlphaFold2,” Communications Biology 8, no. 1 (2025): 373, 10.1038/s42003-025-07791-9.

10. Y. Kalakoti, and B. Wallner, “AFsample3: Generating and Selecting Multiple Conformational States with Alphafold3,” AFsample3: Generating and selecting multiple conformational states with Alphafold3, Bioinformatics 2026, 10.64898/2026.01.16.699904.

11. H. Tian, X. Jiang, F. Trozzi, S. Xiao, E. C. Larson, and P. Tao, “Explore Protein Conformational Space With Variational Autoencoder,” Frontiers in Molecular Biosciences 8 (2021): 781635, 10.3389/fmolb.2021.781635.

12. S. Mansoor, M. Baek, H. Park, G. R. Lee, and D. Baker, “Protein Ensemble Generation Through Variational Autoencoder Latent Space Sampling,” Journal of Chemical Theory and Computation 20, no. 7 (2024): 2689–2695, 10.1021/acs.jctc.3c01057.

13. J. Lu, B. Zhong, Z. Zhang, and J. Tang, “Str2Str: A Score-Based Framework for Zero-Shot Protein Conformation Sampling,” Str2Str: A Score-based Framework for Zero-shot Protein Conformation Sampling, arXiv 2024, 10.48550/arXiv.2306.03117.

14. S. Zheng, J. He, C. Liu, et al., “Predicting Equilibrium Distributions for Molecular Systems with Deep Learning,” Nature Machine Intelligence 6, no. 5 (2024): 558–567, 10.1038/s42256-024-00837-3.

15. B. Jing, B. Berger, and T. Jaakkola, “AlphaFold Meets Flow Matching for Generating Protein Ensembles,” AlphaFold Meets Flow Matching for Generating Protein Ensembles, arXiv 2024, 10.48550/arXiv.2402.04845.

16. J. Jumper, R. Evans, A. Pritzel, et al., “Highly Accurate Protein Structure Prediction with AlphaFold,” Nature 596, no. 7873 (2021): 583–589, 10.1038/s41586-021-03819-2.

17. Z. Lin, H. Akin, R. Rao, et al., “Evolutionary-Scale Prediction of Atomic-Level Protein Structure with a Language Model,” Science 379, no. 6637 (2023): 1123–1130, 10.1126/science.ade2574.

18. S. Lewis, T. Hempel, J. Jiménez-Luna, et al., “Scalable Emulation of Protein Equilibrium Ensembles with Generative Deep Learning,” Science 389, no. 6761 (2025): eadv9817, 10.1126/science.adv9817.

19. M. Invernizzi, S. Bottaro, J. O. Streit, et al., “Advancing Protein Ensemble Predictions Across the Order–Disorder Continuum,” Advancing Protein Ensemble Predictions Across the Order–Disorder Continuum, Biophysics 2025, 10.1101/2025.10.18.680935.

20. Y. Zhang, Y. Liu, Z. Ma, M. Li, C. Xu, and H. Gong, “Improving Diffusion-Based Protein Backbone Generation with Global-Geometry-Aware Latent Encoding,” Nature Machine Intelligence 7, no. 7 (2025): 1104–1118, 10.1038/s42256-025-01059-x.

21. J. Yim, A. Campbell, A. Y. K. Foong, et al., “Fast Protein Backbone Generation with SE(3) Flow Matching,” Fast protein backbone generation with SE(3) flow matching, arXiv 2023, 10.48550/arXiv.2310.05297.

22. G. Janson, G. Valdes-Garcia, L. Heo, and M. Feig, “Direct Generation of Protein Conformational Ensembles via Machine Learning,” Nature Communications 14, no. 1 (2023): 774, 10.1038/s41467-023-36443-x.

23. O. M. Dunne, and O. R. Davies, “A Molecular Model for Self-Assembly of the Synaptonemal Complex Protein SYCE3,” Journal of Biological Chemistry 294, no. 23 (2019): 9260–9275, 10.1074/jbc.RA119.008404.

24. L. Rheinemann, T. Thompson, G. Mercenne, et al., “Interactions between AMOT PPxY Motifs and NEDD4L WW Domains Function in HIV-1 Release,” Journal of Biological Chemistry 297, no. 2 (2021): 100975, 10.1016/j.jbc.2021.100975.

25. F. Wilcoxon, “Individual Comparisons by Ranking Methods,” Biometrics Bulletin 1, no. 6 (1945): 80, 10.2307/3001968.

26. J. Li, K. C. Bennett, Y. Liu, M. V. Martin, and T. Head-Gordon, “Accurate Prediction of Chemical Shifts for Aqueous Protein Structure on ‘Real World’ Data,” Chemical Science 11, no. 12 (2020): 3180–3191, 10.1039/C9SC06561J.

27. S. Grudinin, M. Garkavenko, and A. Kazennov, “Pepsi-SAXS: An Adaptive Method for Rapid and Accurate Computation of Small-Angle X-Ray Scattering Profiles,” Acta Crystallographica Section D Structural Biology 73, no. 5 (2017): 449–464, 10.1107/S2059798317005745.

28. T. N. Nguyen, and D. E. Wemmer, “Letter to the Editor: Complete Resonance Assignments for the Nudix Hydrolase DR2356 of Deinococcus Radiodurans,” Journal of Biomolecular NMR 27, no. 2 (2003): 181–182, 10.1023/A:1024934325331.

29. T. Hošek, E. O. Calçada, M. O. Nogueira, et al., “Structural and Dynamic Characterization of the Molecular Hub Early Region 1A (E1A) from Human Adenovirus,” Chemistry – A European Journal 22, no. 37 (2016): 13010–13013, 10.1002/chem.201602510.

30. P. Upla, S. J. Kim, P. Sampathkumar, et al., “Molecular Architecture of the Major Membrane Ring Component of the Nuclear Pore Complex,” Structure 25, no. 3 (2017): 434–445, 10.1016/j.str.2017.01.006.

31. E. Aragón, Q. Wang, Y. Zou, et al., “Structural Basis for Distinct Roles of SMAD2 and SMAD3 in FOXH1 Pioneer-Directed TGF-β Signaling,” Genes & Development 33, nos. 21–22 (2019): 1506–1524, 10.1101/gad.330837.119.

32. Y. Zhang, and J. Skolnick, “Scoring Function for Automated Assessment of Protein Structure Template Quality,” Proteins: Structure, Function, and Bioinformatics 57, no. 4 (2004): 702–710, 10.1002/prot.20264.

33. P. C. Whitford, O. Miyashita, Y. Levy, and J. N. Onuchic, “Conformational Transitions of Adenylate Kinase: Switching by Cracking,” Journal of Molecular Biology 366, no. 5 (2007): 1661–1671, 10.1016/j.jmb.2006.11.085.

34. J. B. Thoden, C. Z. Blanchard, H. M. Holden, and G. L. Waldrop, “Movement of the Biotin Carboxylase B-Domain as a Result of ATP Binding,” Journal of Biological Chemistry 275, no. 21 (2000): 16183–16190, 10.1074/jbc.275.21.16183.

35. G. Zhang, J. Dai, L. Wang, D. Dunaway-Mariano, L. W. Tremblay, and K. N. Allen, “Catalytic Cycling in β-Phosphoglucomutase: A Kinetic and Structural Analysis,” Biochemistry 44, no. 27 (2005): 9404–9416, 10.1021/bi050558p.

36. D.-A. Silva, G. R. Bowman, A. Sosa-Peinado, and X. Huang, “A Role for Both Conformational Selection and Induced Fit in Ligand Binding by the LAO Protein,” PLoS Computational Biology 7, no. 5 (2011): e1002054, 10.1371/journal.pcbi.1002054.

37. L. McInnes, J. Healy, and J. Melville, “UMAP: Uniform Manifold Approximation and Projection for Dimension Reduction,” UMAP: Uniform Manifold Approximation and Projection for Dimension Reduction, arXiv 2020, 10.48550/arXiv.1802.03426.

38. M. Steinegger, and J. Söding, “MMseqs2 Enables Sensitive Protein Sequence Searching for the Analysis of Massive Data Sets,” Nature Biotechnology 35, no. 11 (2017): 1026–1028, 10.1038/nbt.3988.

39. J. C. Hoch, K. Baskaran, H. Burr, et al., “Biological Magnetic Resonance Data Bank,” Nucleic Acids Research 51, no. D1 (2023): D368–D376, 10.1093/nar/gkac1050.

40. A. G. Kikhney, C. R. Borges, D. S. Molodenskiy, C. M. Jeffries, and D. I. Svergun, “SASBDB: Towards an Automatically Curated and Validated Repository for Biological Scattering Data,” Protein Science 29, no. 1 (2020): 66–75, 10.1002/pro.3731.

41. M. Mirdita, K. Schütze, Y. Moriwaki, L. Heo, S. Ovchinnikov, and M. Steinegger, “ColabFold: Making Protein Folding Accessible to All,” Nature Methods 19, no. 6 (2022): 679–682, 10.1038/s41592-022-01488-1.

42. B. E. Suzek, Y. Wang, H. Huang, P. B. McGarvey, C. H. Wu, and the UniProt Consortium, “UniRef Clusters: A Comprehensive and Scalable Alternative for Improving Sequence Similarity Searches,” Bioinformatics 31, no. 6 (2015): 926–932, 10.1093/bioinformatics/btu739.

43. M. Steinegger, M. Mirdita, and J. Söding, “Protein-Level Assembly Increases Protein Sequence Recovery from Metagenomic Samples Manyfold,” Nature Methods 16, no. 7 (2019): 603–606, 10.1038/s41592-019-0437-4.

44. L. Richardson, B. Allen, G. Baldi, et al., “MGnify: The Microbiome Sequence Data Analysis Resource in 2023,” Nucleic Acids Research 51, no. D1 (2023): D753–D759, 10.1093/nar/gkac1080.

45. S. Chaudhury, S. Lyskov, and J. J. Gray, “PyRosetta: A Script-Based Interface for Implementing Molecular Modeling Algorithms Using Rosetta,” Bioinformatics 26, no. 5 (2010): 689–691, 10.1093/bioinformatics/btq007.

46. V. G. Satorras, E. Hoogeboom, and M. Welling, “E(n) Equivariant Graph Neural Networks,” E(n) Equivariant Graph Neural Networks, arXiv 2021, 10.48550/ARXIV.2102.09844.

47. A. Vaswani, N. Shazeer, N. Parmar, et al., “Attention Is All You Need,” Attention Is All You Need, arXiv 2023, 10.48550/arXiv.1706.03762.

48. D. P. Kingma, and M. Welling, “Auto-Encoding Variational Bayes,” Auto-Encoding Variational Bayes, arXiv 2022, 10.48550/arXiv.1312.6114.

49. R. T. Q. Chen, Y. Rubanova, J. Bettencourt, and D. Duvenaud, “Neural Ordinary Differential Equations,” Neural Ordinary Differential Equations, arXiv 2018, 10.48550/ARXIV.1806.07366.

50. Y. Lipman, R. T. Q. Chen, H. Ben-Hamu, M. Nickel, and M. Le, “Flow Matching for Generative Modeling,” Flow Matching for Generative Modeling, arXiv 2023, 10.48550/arXiv.2210.02747.

