## Supplementary Information for "TopoFlow: Evolutionarily Conditioned Flow Matching for Protein Conformational Ensemble Generation"

**This file includes:**

- Supplementary Notes S1 to S4
- Supplementary Figures S1 to S5
- Supplementary Tables S1 to S7
- Supplementary Algorithms S1 to S3
- Reference

### Supplementary Notes

#### Note S1. Implementation of the MSA Clustering Strategy

For each target protein, an initial multiple sequence alignment (MSA) was generated using ColabFold [1] with MMseqs2 searches. To construct multiple evolutionary conditions, the initial MSA was further processed and partitioned using a sequence-clustering strategy [2]. Before clustering, the query sequence was separated from the MSA and retained for subsequent addition to each clustered MSA. Lowercase insertion characters were removed from the non-query sequences, leaving aligned uppercase amino-acid characters and gap characters. The processed non-query sequences were then one-hot encoded across the aligned MSA positions using an alphabet consisting of the 20 standard amino acids and the gap character, yielding fixed-length representations for subsequent clustering. Five MSA conditions were ultimately constructed from the resulting sequence groups and used for downstream Evoformer representation extraction.

We first applied DBSCAN [2] to the processed non-query sequences to identify fine-grained evolutionary subgroups. The minimum number of samples required to form a dense region was set to 3. To determine the neighborhood radius $\varepsilon$, candidate values ranging from 3.0 to 20.0 in increments of 0.5 were evaluated on a randomly sampled 25% subset of the MSA sequences. The optimal radius was selected as

$$\epsilon^* = \arg\max_{\epsilon \in \mathcal{E}} N_{\text{cluster}}(\varepsilon) \tag{1}$$

where $\mathcal{E} = \{3.0, 3.5, \ldots, 20.0\}$, and $N_{\text{cluster}}(\varepsilon)$ denotes the number of non-noises DBSCAN clusters obtained using radius $\varepsilon$. The selected radius $\varepsilon^*$was subsequently applied to the complete set of processed non-query sequences. Sequences assigned to the DBSCAN noise label were excluded from the subsequent cluster-merging procedure.

Each retained DBSCAN cluster was represented by a consensus sequence computed from the aligned sequences assigned to that cluster. For DBSCAN cluster $C_k$, the consensus residue at aligned position $j$ was defined as

$$c_{k,j} = \arg\max_{a \in \mathcal{A}} \sum_{s_i \in C_k} \mathbf{1}\left( s_j = a \right) \tag{2}$$

where $\mathcal{A}$ denotes the alphabet consisting of the 20 standard amino acids and the gap character, $s_j$ denotes the character at aligned position $j$ in sequence $s$, and $\mathbf{1}(\cdot)$ is the indicator function. The resulting consensus sequences were one-hot encoded and used as cluster-level representations. K-means clustering was then applied to these representations to merge the fine-grained DBSCAN clusters into five higher-level MSA groups.

The resulting consensus sequences were one-hot encoded and used as representative feature vectors for the corresponding DBSCAN clusters. K-means clustering with $k = 5$ was then applied to these cluster-level representations, assigning each DBSCAN cluster to one of five higher-level groups according to the similarity of their consensus sequences. All original MSA sequences belonging to DBSCAN clusters assigned to the same k-means group were subsequently combined to construct the corresponding clustered MSA. In this way, the variable number of fine-grained DBSCAN clusters was consolidated into five MSA groups for downstream conditioning.

**Note S2. Implementation of the VAE Encoder and Latent Representation Learning**

As shown in Supplementary Figure S1, the VAE encoder converts structure-derived features into structural latent variables. The input features comprise Rosetta energy terms [3] and residue-pair distance and orientation maps, which provide complementary energetic and geometric information for each protein conformation. These features are first processed by six equivariant graph neural network (EGNN) layers to integrate local geometric context and residue-pair structural information. The resulting residue embeddings are then refined by a Transformer encoder. Finally, two separate multilayer perceptron (MLP) heads predict the posterior mean and log standard deviation for each residue, thereby parameterizing a residue-wise diagonal Gaussian posterior. For a protein of length $L$, the latent variables are sampled using the reparameterization trick:

$$z_i = \mu_{\phi,i} + \tau_z \sigma_{\phi,i} \odot \epsilon_i, \epsilon_i \sim \mathcal{N}(0, I) \tag{3}$$

where $z_i \in \mathbb{R}^{32}$ denotes the structural latent variable for residue $i$, $\mu_{\phi,i}$ and $\sigma_{\phi,i}$ denote the corresponding posterior mean and standard deviation, respectively, $\tau_z$ controls the stochasticity of latent sampling, and $\odot$ denotes element-wise multiplication. This formulation enables stochastic latent-variable sampling while preserving differentiability during training.

To encourage the latent representation to encode information related to conformational flexibility, residue-level RMSF values derived from the ATLAS MD trajectories are used as supervised labels during VAE pretraining. RMSF values are standardized separately within each target protein. A temporary RMSF prediction head maps the sampled residue-wise latent variables $z_i$to standardized RMSF predictions. The RMSF prediction loss is computed over valid residues as

$$\mathcal{L}_{\mathrm{RMSF}} = \frac{\sum_{i=1}^{L} m_i \,\mathrm{SmoothL1}(\hat{y}_i^{\mathrm{pred}}, \hat{y}_i)}{\sum_{i=1}^{L} m_i} \tag{4}$$

where $m_i$ is the valid-residue mask, $\hat{y}_i^{\mathrm{pred}}$ is the predicted standardized RMSF value, and $\hat{y}_i$ is the corresponding target value. A Kullback–Leibler (KL) regularization term constrains the residue-wise posterior toward a standard normal prior [4]:

$$\mathcal{L}_{\mathrm{KL}} = D_{\mathrm{KL}}\left[q_\phi(z \mid x) \parallel \mathcal{N}(\mathbf{0}, \mathbf{I})\right] \tag{5}$$

The total VAE pretraining objective is

$$\mathcal{L}_{\mathrm{VAE}} = \lambda_{\mathrm{RMSF}} \mathcal{L}_{\mathrm{RMSF}} + \beta_{\mathrm{KL}} \mathcal{L}_{\mathrm{KL}} \tag{6}$$

where $\lambda_{\mathrm{RMSF}}$ and $\beta_{\mathrm{KL}}$ control the contributions of the RMSF prediction and KL regularization terms, respectively. The VAE encoder and the temporary RMSF prediction head are jointly optimized during pretraining. After pretraining, the RMSF head is discarded, and the pretrained VAE encoder is retained for subsequent TopoFlow training.

**Note S3. Implementation of Conditional Flow Matching**

TopoFlow performs conditional flow matching on residue-level rigid frames. For a protein of length $L$, the frame of residue $i$ is represented by a translation $r_i \in \mathbb{R}^3$ and a rotation $R_i \in \mathrm{SO}(3)$. Given the initial frame $T_{0,i} = (R_{0,i}, r_{0,i})$ and the clean endpoint frame $T_{1,i} = (R_{1,i}, r_{1,i})$, the intermediate translational and rotational states are constructed by linear interpolation in $\mathbb{R}^3$ and geodesic interpolation on $\mathrm{SO}(3)$, respectively [5]:

$$r_{t,i} = (1 - t) r_{0,i} + t r_{1,i} \tag{7}$$

$$R_{t,i} = R_{0,i} \exp\left(t \log\left(R_{0,i}^T R_{1,i}\right)\right) \tag{8}$$

where $\exp(\cdot)$ and $\log(\cdot)$ denote the exponential and logarithm maps on $\mathrm{SO}(3)$, respectively. Given the intermediate frame $T_{t,i} = (R_{t,i}, r_{t,i})$ and the time variable $t$, the network predicts the corresponding clean endpoint frame $(\hat{R}_{1,i}, \hat{r}_{1,i})$. The predicted translational vector field is computed as [5]

$$v_{\theta,i}^r = \frac{\hat{r}_{1,i} - r_{t,i}}{1 - t} \tag{9}$$

whereas the rotational vector field is analogously derived from the remaining relative rotation between $R_{t,i}$ and $\hat{R}_{1,i}$ on $\mathrm{SO}(3)$.

The flow-matching objective is optimized together with auxiliary structural reconstruction and latent regularization losses. Since TopoFlow predicts clean endpoint frames rather than directly parameterizing the instantaneous vector field, the translation endpoint loss is defined between the predicted and target clean translations:

$$\mathcal{L}_{trans} = \frac{1}{L} \sum_{i=1}^{L} m_i \left\| \hat{r}_{1,i} - r_{1,i} \right\|_2^2 \tag{10}$$

where $L$ denotes the protein length, $i$ indexes residues, $m_i$represents the residue validity mask, and $\hat{r}_{1,i}$ and $r_{1,i}$ denote the predicted and target endpoint translations, respectively. For rotational dynamics, the loss compares the logarithmic rotation vectors derived from the predicted and target endpoint rotations:

$$\mathcal{L}_{rot} = \frac{1}{L} \sum_{i=1}^{L} m_i \left\| \log\left(R_{t,i}^T R_{1,i}\right) - \log\left(R_{t,i}^T \hat{R}_{1,i}\right) \right\|_2^2 \tag{11}$$

where $R_{t,i}$ denotes the intermediate rotation state, while $R_{1,i}$ and $\hat{R}_{1,i}$ represent the target and predicted endpoint rotations, respectively. The logarithm map projects relative rotations from $SO(3)$ onto the corresponding rotation vectors. The latent representation is regularized by minimizing the divergence between the structural posterior and the sequence-conditioned prior:

$$\mathcal{L}_{KL} = D_{KL}\left(q_\phi(z \mid x_{struct}) \parallel p_\psi(z \mid a_{type})\right) \tag{12}$$

where $q_{\phi}(z \mid x_{struct})$ denotes the posterior distribution encoded from structural features, and $p_{\psi}(z \mid a_{type})$ represents the sequence-conditioned latent prior. Finally, the overall training objective is:

$$\mathcal{L}_{total} = \mathcal{L}_{trans} + \mathcal{L}_{rot} + \beta_{KL}\mathcal{L}_{KL} \tag{13}$$

The complete training and inference procedures are summarized in Supplementary Algorithms S2 and S3, respectively.

**Note S4. Evaluation metrics**

Following Str2Str [6], distributional agreement between generated ensembles and reference MD ensembles was evaluated using JS-PwD, JS-Rg, and JS-TIC. In all three metrics, distributions were represented by 50-bin histograms, with a pseudo count of $\epsilon = 10^{-6}$ added to each bin. Jensen–Shannon distance was calculated as [6]

$$d_{\mathrm{JS}}(P, Q) = \sqrt{\frac{1}{2}D_{\mathrm{KL}}(P \parallel M) + \frac{1}{2}D_{\mathrm{KL}}(Q \parallel M)}, M = \frac{P + Q}{2} \tag{14}$$

**(1) Jensen–Shannon distance for pairwise CA distances (JS-PwD).** For each residue pair $(i, j)$ satisfying $j - i \geq 3$, the CA–CA distance distribution was calculated across all conformations. Generated and reference distributions were constructed using the distance range observed in the reference MD ensemble. JS-PwD was obtained by averaging the Jensen–Shannon distances over all retained residue pairs.

**(2) Jensen–Shannon distance for the radius of gyration (JS-Rg).** For each conformation, the radius of gyration was calculated from the $\mathrm{C}_\alpha$ coordinates as [6]

$$R_g = \sqrt{\frac{1}{N}\sum_{n=1}^{N} \parallel x_n - \bar{x} \parallel_2^2}, \bar{x} = \frac{1}{N}\sum_{n=1}^{N} x_n \tag{15}$$

The generated and reference $R_g$ distributions were then compared using the Jensen–Shannon distance.

**(3) Jensen–Shannon distance for the top two TICs (JS-TIC).** Pairwise $\mathrm{C}_\alpha$ distances were used as input features for time-lagged independent component analysis [7]. TICA was fitted on the reference MD trajectory with a lag time of 20 and two output dimensions, and the fitted model was used to project both ensembles. Jensen–Shannon distances were computed separately for the first two TIC dimensions and averaged to obtain JS-TIC.

Following AlphaFlow [8], ensemble-level agreement between generated ensembles and reference MD ensembles was further evaluated using RMSF Pearson correlation, weak contact agreement, and transient contact agreement. Contact-based metrics were computed using the definitions and thresholds adopted in AlphaFlow.

**(4) RMSF Pearson correlation.** Residue-level flexibility was evaluated using the Pearson correlation between the RMSF profiles of the generated and reference MD ensembles. Before RMSF calculation, all conformations in both ensembles were globally aligned to the static structure used to initialize the corresponding MD simulation. For residue $i$, the RMSF over an ensemble of $K$ conformations was computed as[8]

$$\mathrm{RMSF}_i = \sqrt{\frac{1}{K}\sum_{k=1}^{K} \| x_i^{(k)} - \bar{x}_i \|_2^2} \tag{16}$$

$$\bar{x}_i = \frac{1}{K}\sum_{k=1}^{K} x_i^{(k)} \tag{17}$$

Here, $x_i^{(k)}$ denotes the $\mathrm{C}_\alpha$ coordinate of residue $i$ in conformation $k$, and $\bar{x}_i$ is its ensemble-averaged coordinate. Pearson correlation was calculated between the generated and reference RMSF profiles for each protein and then summarized across the test set. Higher values indicate better agreement in residue-level flexibility.

**(5) Weak contact agreement.** A residue pair was considered to be in contact when its $\mathrm{C}_\alpha$–$\mathrm{C}_\alpha$ distance was less than 8 Å. Weak contacts were defined as residue pairs that were in contact in the static reference structure but dissociated in more than 10% of the ensemble conformations. Agreement between the weak-contact sets obtained from the generated and reference MD ensembles was quantified using Jaccard similarity.

**(6) Transient contact agreement.** Transient contacts were defined as residue pairs that were not in contact in the static reference structure but formed contacts in more than 10% of the ensemble conformations. Agreement between the generated and reference transient-contact sets was also quantified using Jaccard similarity. For a generated contact set $C_{\mathrm{gen}}$ and a reference contact set $C_{\mathrm{ref}}$, Jaccard similarity was calculated as [8]

$$J(C_{\mathrm{gen}}, C_{\mathrm{ref}}) = \frac{| C_{\mathrm{gen}} \cap C_{\mathrm{ref}} |}{| C_{\mathrm{gen}} \cup C_{\mathrm{ref}} |} \tag{18}$$

Following PeptoneBench [9], agreement between generated ensembles and experimental observables was evaluated using uncertainty-normalized RMSE before and after maximum-entropy ensemble reweighting. Performance across the order–disorder continuum was summarized using the area under a LOWESS-smoothed RMSE curve.

**(7) PeptoneBench uncertainty-normalized RMSE.** For each generated ensemble, NMR chemical shifts were predicted using UCBShift [10], and SAXS profiles were computed using Pepsi-SAXS [11]. For $M$ experimental observables, the uncertainty-normalized RMSE was calculated as [9]

$$\mathrm{RMSE} = \sqrt{\frac{1}{M}\sum_{j=1}^{M}\left(\frac{\sum_{i=1}^{N} w_i\, o_j^{\mathrm{gen}}(x_i) - o_j^{\mathrm{exp}}}{\sigma_j}\right)^2} \tag{19}$$

where $N$ is the number of conformations, $w_i$ is the weight of conformation $i$, $o_j^{\mathrm{gen}}(x_i)$ is the forward-model prediction for observable $j$, $o_j^{\mathrm{exp}}$ is the corresponding experimental value, and $\sigma_j$ is its uncertainty. For chemical shifts, disorder-dependent uncertainties interpolating between the UCBShift forward-model uncertainties and POTENCI [12] random-coil prediction uncertainties were used. For SAXS profiles, experimental uncertainties were rescaled following the PeptoneBench procedure, and the ensemble-averaged calculated profile was scaled to the experimental intensity before RMSE calculation. Lower RMSE values indicate better agreement with experimental measurements.

**(8) Maximum-entropy ensemble reweighting.** For unreweighted ensembles, uniform weights $w_i = 1/N$ were assigned. For reweighted ensembles, the conformational weights were optimized by minimizing [9]

$$\mathcal{L}_\theta(\mathbf{w}) = \frac{M}{2}\mathrm{RMSE}^2 + \theta \sum_{i=1}^{N} w_i \log w_i, \sum_{i=1}^{N} w_i = 1 \tag{20}$$

where $\theta$ controls the trade-off between agreement with experimental data and preservation of ensemble diversity. Diversity after reweighting was monitored using the Kish effective sample size (ESS):

$$\mathrm{ESS} = \frac{1}{\sum_{i=1}^{N} w_i^2} \tag{21}$$

Following PeptoneBench, $\theta$ was selected such that the ESS was approximately $10\%$ of the original ensemble size. This constraint prevents the reweighted average from being dominated by only a few conformations.

**(9) LOWESS-AUC aggregation.** For each protein, normalized RMSE was analyzed as a function of disorder content quantified by the mean G-score. LOWESS regression was applied using a smoothing fraction of $0.5$ and one robustifying iteration. The aggregate performance was calculated as the area under the LOWESS curve [9]:

$$\mathrm{LOWESS} - \mathrm{AUC} = \int_0^1 f_{\mathrm{LOWESS}}(g)\, dg \tag{22}$$

where $g$ denotes the mean protein G-score and $f_{\mathrm{LOWESS}}(g)$ is the smoothed normalized RMSE. Uncertainty in the LOWESS curve was estimated using 200 bootstrap resamples, from which 95% confidence intervals were obtained. Lower LOWESS-AUC values indicate better overall agreement with experimental observables across the order–disorder continuum.

**(10) Structure recovery for domain motions.** Following BioEmu [13], recovery of experimentally resolved domain-motion states was primarily evaluated using global CA RMSD. Each generated conformation was globally aligned to each reference conformation using the common CA atoms. For two aligned structures $X$ and $Y$ containing $N$aligned CA atoms, RMSD was calculated as

$$\mathrm{RMSD}(X,Y) = \sqrt{\frac{1}{N}\sum_{n=1}^{N} \| \mathrm{Align}(X)_n - Y_n \|_2^2} \quad (23)$$

TM-score [14] was additionally reported as a complementary length-normalized measure of global structural similarity:

$$\mathrm{TM}(X,Y) = \max_{\mathrm{Align}} \frac{1}{L}\sum_{i=1}^{L} \frac{1}{1 + [d_i/d_0(L)]^2} \quad (24)$$

For each reference conformation, the minimum global CA RMSD and maximum TM-score across all generated conformations were reported. A reference conformation was considered recovered if at least $0.1\%$ of the generated conformations achieved a global CA RMSD of 3Å or less. Lower minimum RMSD values, higher maximum TM-scores, and higher recovery rates indicate better recovery of alternative conformational states.

#### Supplementary Figures

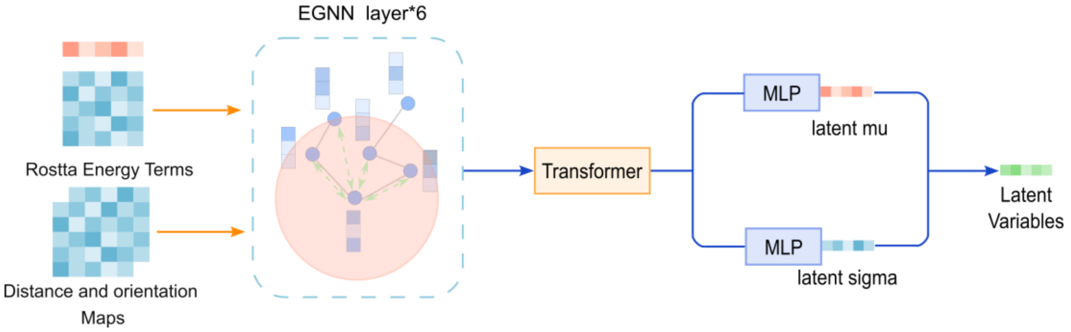


**Fig. S1** Architecture of the VAE encoder. Rosetta energy terms and residue-pair distance and orientation maps are processed by six EGNN layers followed by a Transformer encoder. Two MLP heads predict the residue-wise posterior mean $\mu$ and standard deviation $\sigma$, which parameterize a diagonal Gaussian posterior for structural latent-variable sampling.

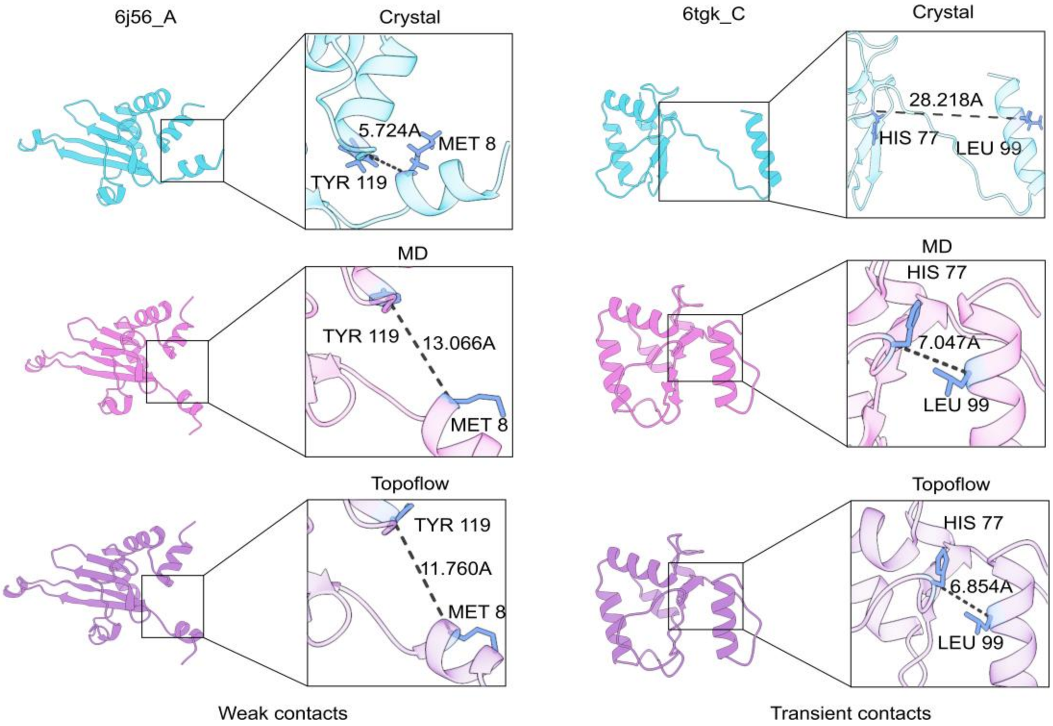


**Fig. S2** Representative weak and transient contacts in 6J56 and 6TGK, showing TopoFlow reproducing weak contact dissociation and transient contact formation.

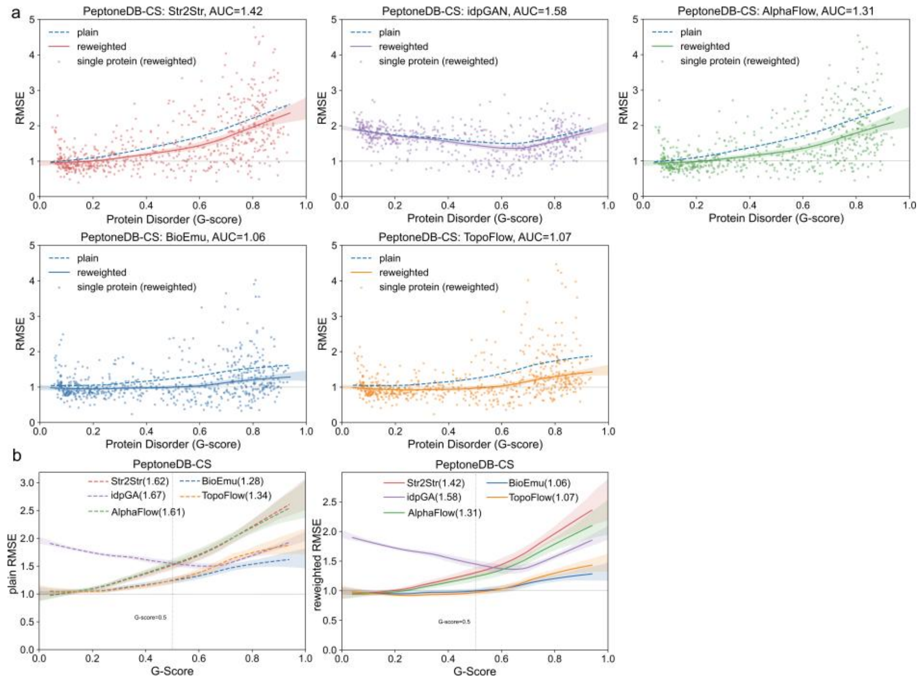


**Fig. S3.** PeptoneDB-CS performance across the protein order–disorder continuum.**a.** Per-protein uncertainty-normalized RMSE values are plotted against mean G-score for Str2Str, idpGAN, AlphaFlow, BioEmu, and TopoFlow. Dashed and solid curves denote LOWESS fits before and after maximum-entropy reweighting, respectively, and the corresponding reweighted LOWESS-AUC is reported for each method.**b.** Comparison of LOWESS-smoothed plain and reweighted RMSE profiles across methods. Numbers in parentheses indicate the corresponding LOWESS-AUC values. The vertical dashed line at $G$-score $= 0.5$ separates the lower- and higher-disorder regimes.

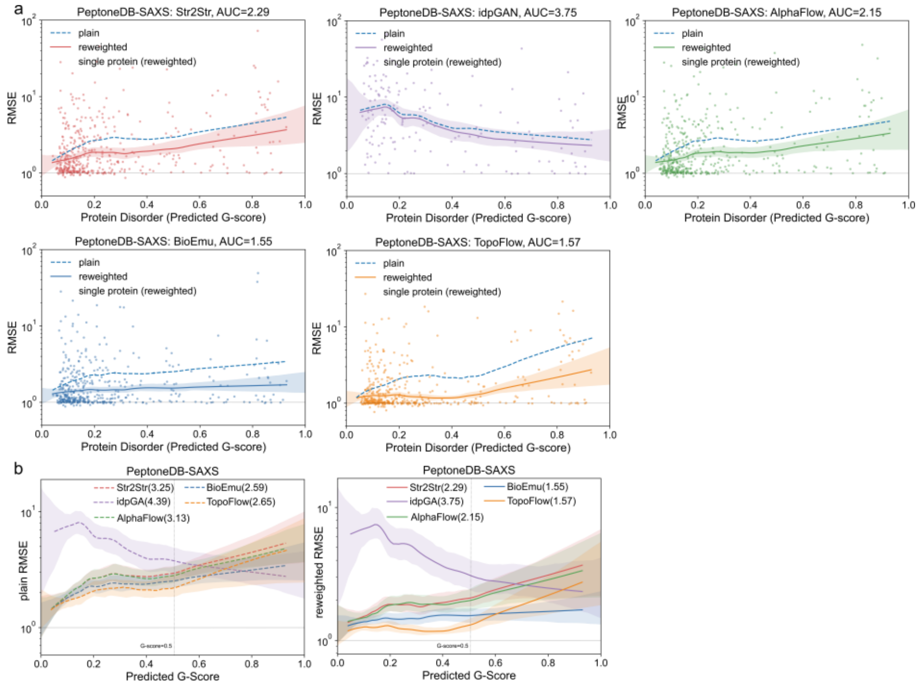


**Fig. S4** PeptoneDB-SAXS performance across the protein order–disorder continuum. **a.**Per-protein uncertainty-normalized RMSE as a function of predicted mean G-score for all methods, with LOWESS fits before and after maximum-entropy reweighting. Reweighted LOWESS-AUC values are shown in each panel. **b.**LOWESS-smoothed plain and reweighted RMSE profiles across methods; values in parentheses denote LOWESS-AUC. The vertical dashed line marks $G$-score $= 0.5$.

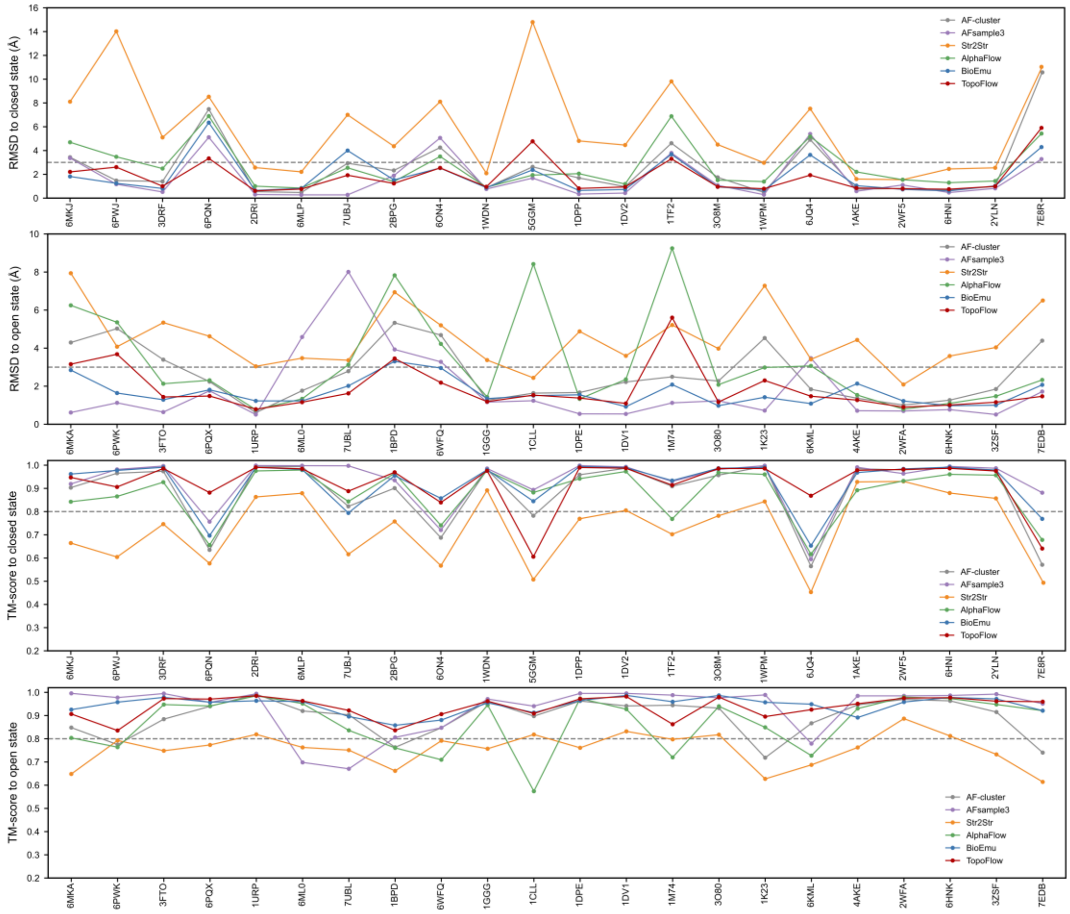


**Fig. S5** Domain motion benchmark per-target structural recovery. RMSD (Å) and TM-score for closed and open reference states are shown for each protein target. TopoFlow generally achieves lower RMSD and higher TM-score compared with baseline methods, indicating effective recovery of both closed and open conformational states across diverse proteins.

### Supplementary Tables

**Table S1.** Performance comparison of TopoFlow and baseline methods on the ATLAS test set. Lower values indicate better performance for JS-PwD, JS-Rg, and JS-TIC, whereas higher values indicate better performance for Jaccard similarity and RMSF correlation.

| Methods | Str2Str | idpGAN | AlphaFlow | BioEmu | TopoFlow |
|---|---|---|---|---|---|
| JS-PwD | 0.437 | 0.401 | 0.372 | 0.354 | 0.316 |
| JS-Rg | 0.426 | 0.416 | 0.368 | 0.346 | 0.326 |
| JS-TIC | 0.429 | 0.402 | 0.390 | 0.374 | 0.345 |
| Weak contact $Ja$ | 0.543 | 0.562 | 0.645 | 0.615 | 0.668 |
| Transient contact $Ja$ | 0.356 | 0.347 | 0.392 | 0.398 | 0.425 |
| RMSF r | 0.768 | 0.783 | 0.829 | 0.812 | 0.854 |

**Table S2.** Performance comparison based on LOWESS-AUC between TopoFlow and baseline methods on the PeptoneDB-CS benchmark before and after maximum entropy reweighting.

| Methods | Str2Str | idpGAN | AlphaFlow | BioEmu | TopoFlow |
|---|---|---|---|---|---|
| AUC_LOWESS(plain) | 1.616 | 1.667 | 1.619 | 1.282 | 1.342 |
| AUC_LOWESS(reweighted) | 2.29 | 1.582 | 1.312 | 1.058 | 1.066 |

**Table S3.** Performance comparison based on LOWESS-AUC between TopoFlow and baseline methods on the PeptoneDB- SAXS benchmark before and after maximum entropy reweighting.

| Methods | Str2Str | IDPGAN | AlphaFlow | BioEmu | TopoFlow |
|---|---|---|---|---|---|
| AUC_LOWESS(plain) | 3.246 | 4.386 | 3.126 | 2.588 | 2.654 |
| AUC_LOWESS(reweighted) | 2.29 | 3.75 | 2.153 | 1.556 | 1.574 |

**Table S4.** Summarizes the domain motion benchmark results, showing RMSD (Å) and TM-score for closed and open reference conformations for each method.

| Methods | Close state | | Open state | |
|---|---|---|---|---|
| | RMSD (Å) | TM-score | RMSD (Å) | TM-score |
| AF-Cluster | 2.54 | 0.89 | 2.63 | 0.89 |
| AFsample3 | 1.73 | 0.93 | 1.77 | 0.93 |
| Str2Str | 5.91 | 0.73 | 4.49 | 0.76 |
| AlphaFlow | 2.72 | 0.88 | 3.25 | 0.87 |
| BioEmu | 1.86 | 0.92 | 1.66 | 0.94 |
| TopoFlow | 1.82 | 0.92 | 1.68 | 0.94 |

**Table S5.** Structural recovery performance of TopoFlow and baseline methods on the domain motion benchmark.

| Methods | Close state | | Open state | |
|---|---|---|---|---|
| | RMSD ≤3 Å (count) | Success rate | RMSD ≤3 Å (count) | Success rate |
| AF-Cluster | 16 | 73% | 15 | 68% |
| AFsample3 | 16 | 73% | 17 | 77% |
| Str2Str | 8 | 36% | 2 | 9% |
| AlphaFlow | 15 | 68% | 14 | 64% |
| BioEmu | 17 | 77% | 21 | 95% |
| TopoFlow | 18 | 82% | 18 | 82% |

**Table S6.** Ablation study of TopoFlow variants on the ATLAS test set using JS-PwD, JS-Rg, JS-TIC, Weak contacts $Ja$, Transient contacts $Ja$, and RMSF r.

| Configuration | TopoFlow | w/o VAE | w/o Cluster | w/o Cluster & VAE |
|---|---|---|---|---|
| JS-PwD | 0.316 | 0.365 | 0.347 | 0.399 |
| JS-Rg | 0.326 | 0.362 | 0.328 | 0.402 |
| JS-TIC | 0.345 | 0.387 | 0.361 | 0.420 |
| Weak contact $Ja$ | 0.668 | 0.631 | 0.621 | 0.591 |
| Transient contact $Ja$ | 0.425 | 0.427 | 0.383 | 0.351 |
| RMSF r | 0.854 | 0.801 | 0.851 | 0.781 |

**Table S7.** Experimentally resolved open and closed conformational states used in the domain motion benchmark.

| Target | Open state PDB ID | Rg (Å) | Closed state PDB ID | Rg (Å) |
|---|---|---|---|---|
| A0A075Q0W3 | 6MKA_A | 33.22 | 6MKJ_A | 32.51 |
| A0A0H3AFX3 | 6PWK_A | 23.59 | 6PWJ_A | 22.10 |
| A2RJ53 | 3FTO_A | 25.45 | 3DRF_A | 23.58 |
| B0F0C5 | 6PQX_A | 24.61 | 6PQN_A | 23.38 |
| P0205 | 1URP_A | 20.19 | 2DRI_A | 19.20 |
| P02911 | 6ML0_A | 19.12 | 6MLP_A | 17.58 |
| P03047 | 7UBL_A | 18.71 | 7UBJ_A | 16.40 |
| P06766 | 1BPD_A | 28.04 | 2BPG_E | 22.63 |
| P0A8W0 | 6WFQ_A | 24.75 | 6ON4_A | 23.65 |
| P0AEQ3 | 1GGG_A | 19.06 | 1WDN_A | 17.59 |
| P0DP23 | 1CLL_A | 21.89 | 5GGM_A | 16.26 |
| P23847 | 1DPE_A | 24.68 | 1DPP_A | 22.77 |
| P24182 | 1DV1_A | 22.89 | 1DV2_A | 21.74 |
| P28366 | 1M74_A | 33.10 | 1TF2_A | 32.63 |
| P33284 | 3O80_A | 24.14 | 3O8M_A | 23.05 |
| P37487 | 1K23_C | 23.28 | 1WPM_B | 18.66 |
| P67701 | 6KML_B | 22.07 | 6JQ4_A | 20.55 |
| P69441 | 4AKE_A | 19.49 | 1AKE_A | 16.39 |
| P71447 | 2WFA_A | 18.43 | 2WF5_A | 17.05 |
| Q18A65 | 6HNK_A | 20.93 | 6HNI_A | 19.31 |
| Q5F9M1 | 3ZSF_A | 19.49 | 2YLN_A | 17.95 |
| Q83VS8 | 7EDB_A | 22.94 | 7E8R_A | 22.25 |

#### Supplementary Algorithms

---

**Algorithm S1.** RMSF-supervised VAE encoder pretraining

---

def VAEPretrainingLoop($\mathcal{D}_{\text{train}}, E_\phi, f_{\text{RMSF}}, \lambda_{\text{RMSF}}, \beta_{\text{KL}}, \tau_z$):

*# Initialize encoder and temporary RMSF head*

1: initialize $E_\phi = \{\text{EGNN}_\phi, \text{Transformer}_\phi, \text{MLP}_{\mu,\phi}, \text{MLP}_{\sigma,\phi}\}$ and $f_{\text{RMSF}}$

*# RMSF-supervised pretraining*

2: for epoch = 1, …, $N_{\text{epoch}}$ do

3:     for $(x_{\text{node}}, x_{\text{edge}}, r, y, m) \in \mathcal{D}_{\text{train}}$ do

4:         $\hat{y} \leftarrow \text{standardize}(y)$ within each protein

5:         $h \leftarrow \text{EGNN}_\phi(x_{\text{node}}, x_{\text{edge}}, r)$

6:         $h \leftarrow \text{Transformer}_\phi(h, m)$

7:         $\mu_\phi, \log\sigma_\phi \leftarrow \text{MLP}_{\mu,\phi}(h), \text{MLP}_{\sigma,\phi}(h)$

8:         $z_i \leftarrow \mu_{\phi,i} + \tau_z \sigma_{\phi,i} \odot \epsilon_i, \quad \epsilon_i \sim \mathcal{N}(0, I)$

9:         $\hat{y}_i^{\text{pred}} \leftarrow f_{\text{RMSF}}(z_i)$

10:        $\mathcal{L}_{\text{RMSF}} \leftarrow \text{MaskedSmoothL1}(\hat{y}^{\text{pred}}, \hat{y}; m)$

11:        $\mathcal{L}_{\text{VAE}} \leftarrow \lambda_{\text{RMSF}} \mathcal{L}_{\text{RMSF}} + \beta_{\text{KL}} D_{\text{KL}}[q_\phi(z \mid x) \parallel \mathcal{N}(0, I)]$

12:        update $E_\phi$ and $f_{\text{RMSF}}$ by minimizing $\mathcal{L}_{\text{VAE}}$

13:    end for

14: end for

15: discard $f_{\text{RMSF}}$ and return pretrained $E_\phi$

---

**Algorithm S2.** TopoFlow training with pretrained VAE initialization

**def** TopoFlowTrainingLoop($\mathcal{D}_{\text{train}}$, $ckpt_{\text{VAE}}$, $\theta$, $\phi$, $\psi$, $\lambda_{\text{aux}}$, $\beta_{\text{KL}}$):

*# Initialize model and load pretrained VAE encoder*

1: $E_{\phi}$ ← load_pretrained_VAE_encoder($ckpt_{\text{VAE}}$)

2: FlowModel ← initialize_flow_model($E_{\phi}$, SequenceLatentPrior)

*# Joint flow-matching training*

3: **for** epoch = 1, …, $N_{\text{epoch}}$ **do**

4: **for** ($x_{\text{struct}}$, aatype, $s_{\text{Evo}}$, $p_{\text{Evo}}$, $T_1$, $m$) ∈ $\mathcal{D}_{\text{train}}$ **do**

5: $T_t$, $t$ ← Interpolant.corrupt($T_1$)

6: $z$, $\mu_{\phi}$, $\log\sigma_{\phi}$ ← $E_{\phi}(x_{\text{struct}})$

7: $\mu_{\psi}$, $\log\sigma_{\psi}$ ← SequenceLatentPrior(aatype, $m$)

8: $s_{\text{cond}}$, $p_{\text{cond}}$ ← Conditional Modulation Module ($s_{\text{Evo}}$, $p_{\text{Evo}}$, $z$, $t$, $T_t$, $m$)

9: **for** block = 1, …, $N_{\text{block}}$ **do**

10: $s_{\text{cond}}$ ← Invariant Point Attention ($s_{\text{cond}}$, $p_{\text{cond}}$, $T_t$, $m$)

11: $s_{\text{cond}}$ ← Transformer Refinement($s_{\text{cond}}$, $m$)

12: $p_{\text{cond}}$ ← EdgeUpdate($p_{\text{cond}}$, $s_{\text{cond}}$, $m$)

13: $T_t$ ← BackboneFrameUpdate($T_t$, $s_{\text{cond}}$)

14: **end for**

15: $\hat{T}_1$← PredictionHeads($s_{\text{cond}}$, $p_{\text{cond}}$, $T_t$)

16: $\mathcal{L}_{\text{total}} \leftarrow \mathcal{L}_{\text{trans}} + \mathcal{L}_{\text{rot}} + \beta_{\text{KL}} D_{\text{KL}}\left[q_{\phi}(z \mid x_{\text{struct}}) \parallel p_{\psi}(z \mid aatype)\right]$

17: update $\theta$, $\phi$ and $\psi$ by minimizing $\mathcal{L}_{\text{total}}$

18: **end for**

19: **end for**

20: return

**Algorithm S3.** TopoFlow inference and ensemble sampling

**def** TopoFlowInferenceLoop($\mathcal{D}_{\text{test}}$, $ckpt_{\text{flow}}$, $N_{\text{sample}}$, $100$):

*# Load trained model and target conditions*

1: FlowModel ← load_checkpoint($ckpt_{\text{flow}}$)

2: **for** target ∈ $\mathcal{D}_{\text{test}}$ **do**

3: aatype, $s_{\text{Evo}}$, $p_{\text{Evo}}$, $m$ ← load_precomputed_features(target)

4: $\mu_\psi$, $\log\sigma_\psi$ ← SequenceLatentPrior(aatype, $m$)

*# Sample latent variables from sequence prior and initialize noisy frames*

5: **for** sample = 1, …, $N_{\text{sample}}$ **do**

6: $z \leftarrow \mu_\psi + \exp(\log\sigma_\psi) \odot \epsilon, \quad \epsilon \sim \mathcal{N}(0, I)$

7: $T_t$ ← initialize_noisy_frames(aatype)

8: **for** $k$ = 1, …, $100$ **do**

9: $s_{\text{cond}}$, $p_{\text{cond}}$ ← Conditional Modulation Module ($s_{\text{Evo}}$, $p_{\text{Evo}}$, $z$, $t_k$, $T_t$, $m$)

10: **for** block = 1, …, $N_{\text{block}}$ **do**

11: $s_{\text{cond}}$ ← Invariant Point Attention ($s_{\text{cond}}$, $p_{\text{cond}}$, $T_t$, $m$)

12: $s_{\text{cond}}$ ← Transformer ($s_{\text{cond}}$, $m$)

13: $p_{\text{cond}}$ ← Edge Update($p_{\text{cond}}$, $s_{\text{cond}}$, $m$)

14: $T_t$ ← Backbone FrameUpdate($T_t$, $s_{\text{cond}}$)

15: **end for**

16: $\hat{T}_1$← PredictionHeads($s_{\text{cond}}$, $p_{\text{cond}}$, $T_t$)

17: $T_t$ ← Reverse Flow Update($T_t$, $\hat{T}_1$, $t_k$, $\Delta t$)

18: **end for**

19: structure ← Frame-to-Structure($T_1$)

20: **end for**

21: **end for**

22: return generated conformational ensemble

### References


1. M. Mirdita, K. Schütze, Y. Moriwaki, L. Heo, S. Ovchinnikov, and M. Steinegger, "ColabFold: Making Protein Folding Accessible to All," *Nature Methods 19*, no. 6 (2022): 679–682, https://doi.org/10.1038/s41592-022-01488-1.

2. H. K. Wayment-Steele, A. Ojoawo, R. Otten, et al., "Predicting Multiple Conformations via Sequence Clustering and AlphaFold2," *Nature 625*, no. 7996 (2024): 832–839, https://doi.org/10.1038/s41586-023-06832-9.

3. S. Chaudhury, S. Lyskov, and J. J. Gray, "PyRosetta: A Script-Based Interface for Implementing Molecular Modeling Algorithms Using Rosetta," *Bioinformatics 26*, no. 5 (2010): 689–691, https://doi.org/10.1093/bioinformatics/btq007.

4. D. P. Kingma, and M. Welling, "Auto-Encoding Variational Bayes," *Auto-Encoding Variational Bayes*, arXiv 2022, https://doi.org/10.48550/arXiv.1312.6114.

5. J. Yim, A. Campbell, A. Y. K. Foong, et al., "Fast Protein Backbone Generation with SE(3) Flow Matching," *Fast protein backbone generation with SE(3) flow matching*, arXiv 2023, https://doi.org/10.48550/arXiv.2310.05297.

6. J. Lu, B. Zhong, Z. Zhang, and J. Tang, "Str2Str: A Score-Based Framework for Zero-Shot Protein Conformation Sampling," *Str2Str: A Score-based Framework for Zero-shot Protein Conformation Sampling*, arXiv 2024, https://doi.org/10.48550/arXiv.2306.03117.

7. C. R. Schwantes, and V. S. Pande, "Modeling Molecular Kinetics with tICA and the Kernel Trick," *Journal of Chemical Theory and Computation 11*, no. 2 (2015): 600–608, https://doi.org/10.1021/ct5007357.

8. B. Jing, B. Berger, and T. Jaakkola, "AlphaFold Meets Flow Matching for Generating Protein Ensembles," *AlphaFold Meets Flow Matching for Generating Protein Ensembles*, arXiv 2024, https://doi.org/10.48550/arXiv.2402.04845.

9. M. Invernizzi, S. Bottaro, J. O. Streit, et al., "Advancing Protein Ensemble Predictions Across the Order–Disorder Continuum," *Advancing Protein Ensemble Predictions Across the Order–Disorder Continuum*, Biophysics 2025, https://doi.org/10.1101/2025.10.18.680935.

10. J. Li, K. C. Bennett, Y. Liu, M. V. Martin, and T. Head-Gordon, "Accurate Prediction of Chemical Shifts for Aqueous Protein Structure on 'Real World' Data," *Chemical Science 11*, no. 12 (2020): 3180–3191, https://doi.org/10.1039/C9SC06561J.

11. S. Grudinin, M. Garkavenko, and A. Kazennov, "*Pepsi-SAXS*: An Adaptive Method for Rapid and Accurate Computation of Small-Angle X-Ray Scattering Profiles," *Acta Crystallographica Section D Structural Biology 73*, no. 5 (2017): 449–464, https://doi.org/10.1107/S2059798317005745.

12. J. T. Nielsen, and F. A. A. Mulder, "POTENCI: Prediction of Temperature, Neighbor and pH-Corrected Chemical Shifts for Intrinsically Disordered Proteins," *Journal of Biomolecular NMR 70*, no. 3 (2018): 141–165, https://doi.org/10.1007/s10858-018-0166-5.

13. S. Lewis, T. Hempel, J. Jiménez-Luna, et al., "Scalable Emulation of Protein Equilibrium Ensembles with Generative Deep Learning," *Science 389*, no. 6761 (2025): eadv9817, https://doi.org/10.1126/science.adv9817.

14. Y. Zhang, and J. Skolnick, "Scoring Function for Automated Assessment of Protein Structure Template Quality," *Proteins: Structure, Function, and Bioinformatics 57*, no. 4 (2004): 702–710, https://doi.org/10.1002/prot.20264.
